# From spots to stripes: Evolution of pigmentation patterns in monkeyflowers via modulation of a reaction-diffusion system and its prepatterns

**DOI:** 10.1101/2025.01.10.632501

**Authors:** Mei Liang, Lee Ringham, Changning Ye, Xu Yan, Nathan Schaumburger, Mikolaj Cieslak, Michael Blinov, Przemyslaw Prusinkiewicz, Yao-Wu Yuan

## Abstract

The reaction-diffusion (RD) system is widely assumed to account for many complex, self-organized pigmentation patterns in natural organisms. However, the specific configurations of such RD networks and how RD systems interact with positional information (i.e., prepatterns) that may specify the initiation conditions for the RD operation remain largely unknown. Here, we introduced a three-substance RD system underlying the formation of repetitive pigment spots and stripes in *Mimulus* flowers. It consists of an R2R3-MYB activator (NEGAN), an R3-MYB inhibitor (RTO), and a coactivator represented by two paralogous bHLH proteins. Through fine-scale genetic analyses, transgenic experiments, and computer simulations, we identified the causal loci contributing to the evolutionary transition from sparsely dispersed spots to longitudinal stripes. Genetic changes at these loci modulate the prepatterns of the activator and coactivator expression and the promoter activities of the inhibitor and one of the coactivator paralogs. Our findings highlight the importance of prepatterns towards a realistic description of RD systems in natural organisms, and reveal the genetic mechanism generating pattern variation through modulation of the kinetics of the RD system and its prepatterns.

## INTRODUCTION

A classical theoretical framework that explains how gene activities could be translated through cell fate specification into tissue-level patterns is the reaction-diffusion (RD) model. This model was initially proposed by Turing^1^ and then independently developed by Meinhardt and Gierer,^2,3^ who provided the key insight of pattern formation through local self-activation and long-range inhibition. The essence of the Meinhardt-Gierer model is an interplay between two antagonistic substances (morphogens): a pattern-inducing activator, which promotes its own production and diffuses slowly, and another fast-diffusing substance, which represses the production of the activator either through its presence (in the activator-inhibitor model) or absence (in the activator–depleted substrate model). Numerous empirical and simulation studies have suggested that RD-based processes are common in nature^4–8^ and underlie a wide range of developmental patterns.^9–16^ The RD model has two appealing features that make it particularly attractive to the studies of pigmentation systems, from fish skins^9,17,18^ and mammalian coats^19^ to butterfly wings^20^ and flower petals.^21–23^ First, the self-organizing RD system has an intrinsic property of generating patterns with a large number of repetitive elements (e.g., zebra stripes and leopard spots) that are difficult to explain by other developmental mechanisms. Second, subtle changes in the reaction and diffusion parameters can lead to dramatic variation in pattern output, thereby providing an elegant mechanistic solution to the rapid evolution of pigmentation patterns observed in many natural organisms.^24–27^ However, despite the intense interest in using the RD model to explain pigment pattern formation, very few presumptive pattern-inducing factors and their inhibitors have been identified to date, let alone the specific evolutionary changes that contribute to pigment pattern variation in nature via modulation of the RD dynamics.

Here we have studied a group of closely related monkeyflower (*Mimulus*) species, including *M. lewisii*, *M. parishii*, and *M. cardinalis*, that produce anthocyanin pigment spots or stripes in their flowers (**Figure 1A**). Each *Mimulus* flower has five petals (two dorsal, two lateral, and one ventral) that are partially united into a corolla tube. The tubular portion of the ventral petal features two raised ridges with dense trichomes, forming the nectar guides, which occur in all *Mimulus* flowers but are particularly pronounced in *M. lewisii* because of the color contrast with the rest of the corolla (**Figure 1A**). Both *M. lewisii* and *M. parishii* produce small, dispersed anthocyanin spots in the nectar guide area, slightly spreading to the lateral petals, whereas *M. cardinalis* produces longitudinal stripes that spread across all five petals (**Figure 1A**).

**Figure 1.**
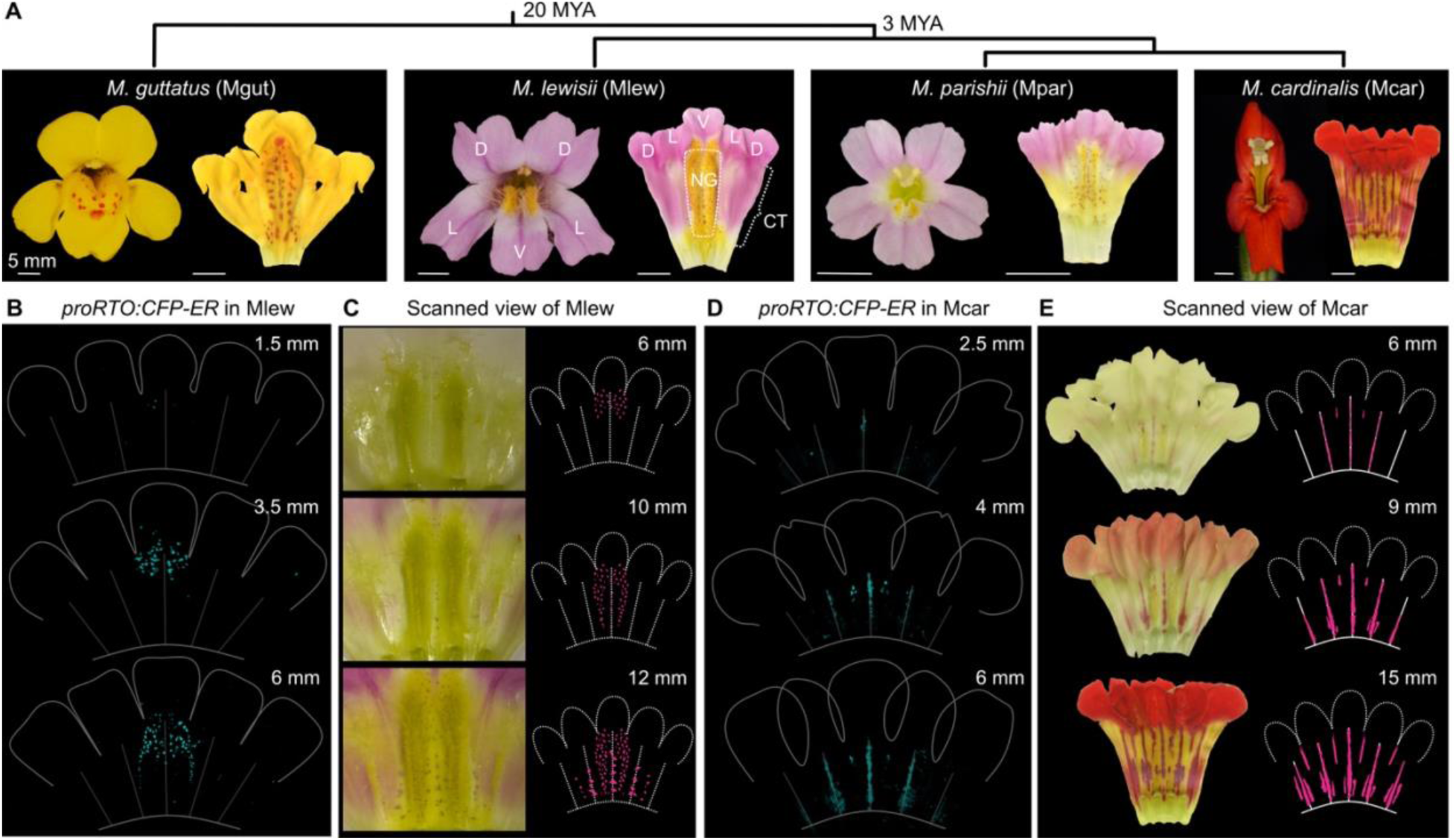
Pigment pattern variation among *Mimulus* species **(A)** Phylogenetic relationships among four *Mimulus* species with estimated divergence time.^28^ Shown for each species are both the front (left) and dissected (right) view of the flower. To produce the dissected view, the corolla was cut open along the junction between the two dorsal petals and then flattened and scanned using a scanner. The dorsal (D), lateral (L), and ventral (V) petals are labeled on the *M. lewisii* flower images, with the yellow nectar guide (NG) area and corolla tube (CT) indicated by the dashed box and bracket, respectively. **(B** and **D)** Confocal fluorescent images showing *RTO* promoter activity at early corolla developmental stages (≤ 6 mm) in *M. lewisii* (B) and *M. cardinalis* (D). **(C** and **E)** Scanned images of later corolla stages (≥ 6 mm) in *M. lewisii* (C) and *M. cardinalis* (E). Corresponding schematic illustrations of the anthocyanin pattern are also shown along the scanned images in (C) and (E).

Previous studies of these *Mimulus* species have identified an anthocyanin-activating R2R3-MYB transcription factor, NEGAN, and an R3-MYB anthocyanin repressor, RTO, that control the formation of dispersed anthocyanin spots in *M. lewisii* and the distantly related *M. guttatus*.^22,29^ This pair of MYB proteins form a local autocatalytic feedback loop and a long-range inhibitory feedback loop, with the following properties: NEGAN activates its own expression and the expression of RTO; RTO exhibits intercellular movement and inhibits NEGAN function along its diffusion path. These properties fulfill the tenets of the classical two-substance, activator-inhibitor RD model.^2–4^ Computer simulations using this model with a random, noisy initiation condition for the activator could recapitulate the pattern changes in mutants and transgenic lines where the expression of the activator (NEGAN) or inhibitor (RTO) was perturbed.^22^

Nevertheless, the above model does not fully capture the biological complexity of the pigmentation patterns in *Mimulus*. First, the spatial distribution of the pigment spots is largely restricted to the nectar guide area in *M. guttatus*, *M. lewisii*, and *M. parishii* (**Figure 1A**), indicating the existence of a prepattern that provides a spatially defined input to the RD process. Indeed, when Ringham et al. incorporated a hypothetical prepattern corresponding to the nectar guide area into the two-substance, activator-inhibitor model and simulated the pattern directly on a geometric model of *M. guttaus* flowers, the output pattern became almost indistinguishable from that of the real flowers.^23^ However, the existence of this prepattern had yet to be experimentally tested and its spatial configuration remained to be determined. Second, the rate of activator production in the basic activator-inhibitor models employed in the previous studies^22,23^ is inversely proportional to the concentration of the inhibitor. This interaction is not consistent with the law of mass action, and thus does not lend itself directly to biochemical implementation.^30,31^ A more plausible biochemical implementation involves a coactivator that is required for the operation of the activator and is removed from the system by the inhibitor. In fact, in the case of anthocyanin pigmentation, it is well known that the R2R3-MYB activator interacts with bHLH and WD40 proteins forming an activation complex, and the R3-MYB inhibitor represses anthocyanin production by competing with the R2R3-MYB for the limited supply of the bHLH coactivators.^32–36^ Yet these coactivators were assumed to remain constant in all petal cells and thus were not included in the RD network in the previous *Mimulus* pigmentation models. Third, while it is easy to imagine how genetic changes can alter the RD parameters and hence spot size or density, it is unclear how to evolve directional stripes (as in *M. cardinalis*) from seemingly randomly dispersed spots (as in *M. lewisii*). Tying the diversity of patterns to evolutionary changes requires a model that more realistically accounts for the operation of the RD system and the putative prepatterns.

In this study, we aim to: (**i**) Determine the prepatterns that control the operation of the RD system during pigment patterning in *Mimulus*; (**ii**) Identify the causal genes contributing to the spot-to-stripe transition from *M. lewisii/parishii* to *M. cardinalis*, with the expectation that some of these genes may encode additional components of the RD network; (**iii**) Develop a biochemically plausible model that incorporates the additional RD network components and their prepatterns; and, (**iv**) Examine how modulation of the dynamics of the RD system and its prepatterns produces evolutionary pattern variation through both experimental manipulation and computer simulation.

## RESULTS

### Developmental dynamics of spot and longitudinal stripe patterning

To unravel the developmental basis of the spot-to-stripe transition, we analyzed the spatiotemporal dynamics of pigment accumulation during flower development in the spotted *M. lewisii* and striped *M. cardinalis*. Because the transcription of anthocyanin biosynthetic genes precedes pigment production and coincides with *RTO* expression (both are activated by NEGAN),^22^ we used the *RTO* promoter reporter as a readout of anthocyanin biosynthesis during early flower developmental stages before the pigments become visible. We first imaged the previously characterized *proRTO:CFP-ER* reporter line in the *M. lewisii* genetic background.^22^ A few CFP signal spots emerged at the top of the nectar guide area (at the junction between the ventral petal lobe and the nectar guides) as early as when the corolla was 1.5 mm long (**Figure 1B**). As the corolla reached 3.5 mm in length, the spotty CFP signals became denser and started covering the top area of the nectar guides. By the 6-mm stage, CFP signals spread along the edges of the nectar guides (**Figure 1B**) and anthocyanin spots became slightly visible to the naked eye (**Figure 1C**). At the 10-mm stage, anthocyanin spots started appearing throughout the nectar guide area, and by the time when the corolla reached 12 mm, sparse anthocyanin spots began to emerge at the bottom part of the lateral petals (**Figure 1C)**.

To image the corresponding processes in *M. cardinalis*, we introduced the *proRTO:CFP-ER* reporter gene into *M. cardinalis* through four rounds of backcrosses. As expected, CFP signals coincide with anthocyanin accumulation (compare the 6-mm stage in **Figures 1D** and **1E**), suggesting that the *proRTO:CFP-ER* reporter can also serve as a readout of anthocyanin biosynthesis in *M. cardinalis*. In contrast to the patterns observed in *M. lewisii*, CFP signals first appeared along the central vein at the petal lobe-nectar guide junction (**Figure 1D**). As the corolla developed, both the CFP signals and the visible pigments emerged sequentially along the central veins of the lateral petals and then the dorsal petals and finally in the interspace between the veins (**Figures 1D** and **1E**). Because both *RTO* and the anthocyanin biosynthetic genes are activated by *NEGAN*,^22^ we speculated that the initiation condition for *NEGAN* expression (i.e. the prepatterns of *NEGAN* expression before the RD process kicks in) might differ between *M. lewisii* and *M. cardinalis*, which may underlie the difference in the initiation location of the CFP signals and the pigment pattern output between the two species.

### Prepatterns of the activator *NEGAN*

To test whether *NEGAN* itself contributes to the pigment pattern variation, we swapped the *NEGAN* alleles between *M. lewisii* and *M. cardinalis* and generated near-isogenic lines (NILs) by introgressing the *NEGAN^L^* allele into *M. cardinalis* and vice versa (**Figure S1A**). In the *M. cardinalis* background, we were only able to obtain a *NEGAN^L/C^*NIL where *NEGAN* is heterozygous, likely due to segregation distortion known to occur in crosses between these species.^37^ However, even at the heterozygous state, the *NEGAN^L^* allele was sufficient to cause an obvious pattern change in *M. cardinalis*: the longitudinal stripes on the ventral and lateral petals became large, disorganized spots (**Figures 2A** and **S1A**). Introgressing both *NEGAN^L^* and *RTO^L^* into *M. cardinalis* resulted in a double NIL (*NEGAN^L/C^ RTO^L/L^*) with further reduced spot size but an overall pattern similar to the *NEGAN^L/C^* NIL (**Figures 2A**). These results support the notion that *NEGAN* has evolved different prepatterns that set its distinct initial expression domain in *M. lewisii* and *M. cardinalis*, and further suggest that these different prepatterns are conferred *in cis* to *NEGAN* (i.e. *cis*-regulatory region of *NEGAN*) instead of through trans-acting factors outside of the introgressed *NEGAN* locus.

**Figure 2.**
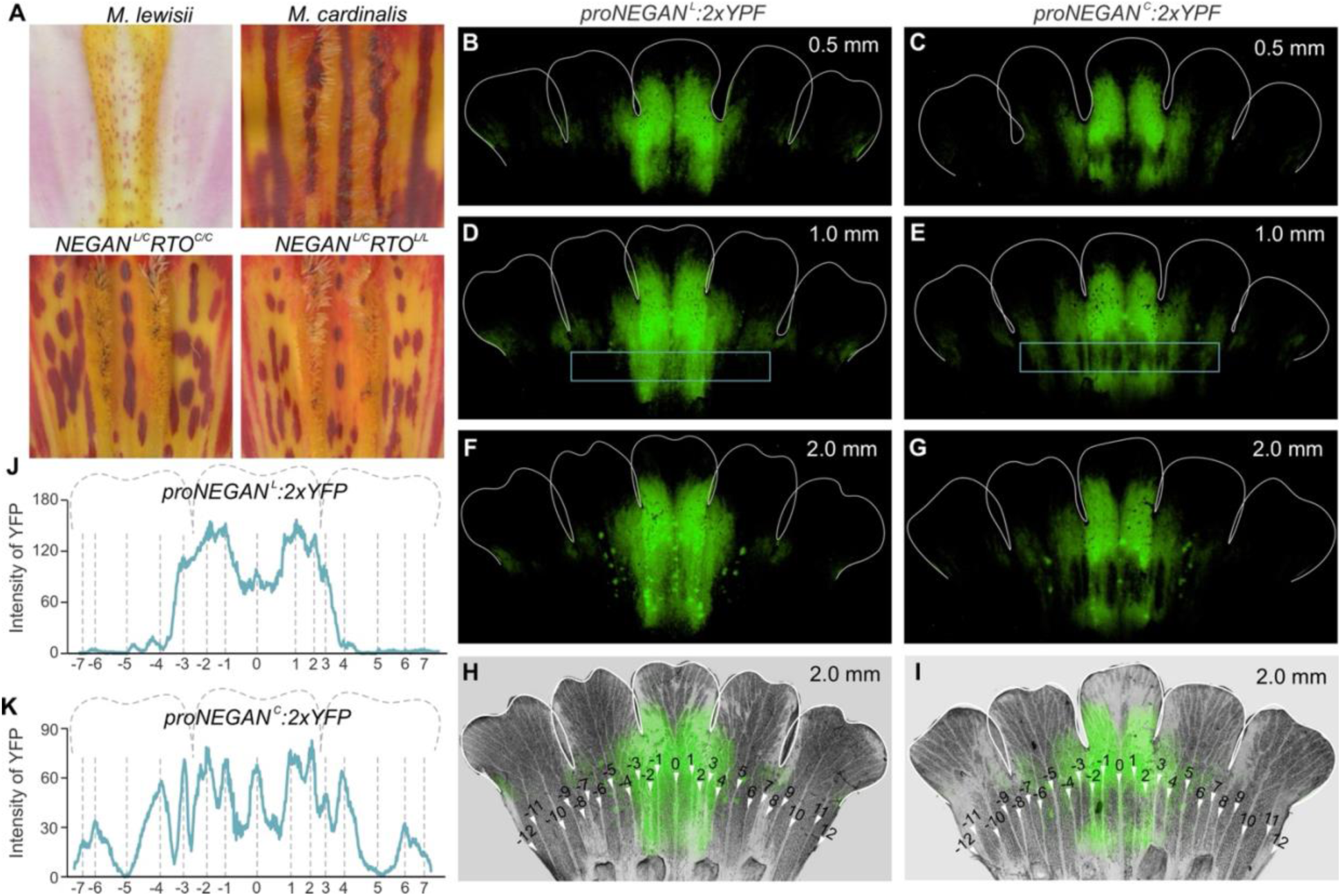
Prepatterns of *NEGAN* expression **(A)** Introgressing the *M. lewisii NEGAN* allele (*NEGAN^L^*) into *M. cardinalis*, even at the heterozygous state, resulted in pattern change from longitudinal stripes to disorganized spots. (**B** to **G**) The reporter line of *M. lewisii NEGAN* promoter (*proNEGAN^L^*) showed a different fluorescence pattern compared to the *M. cardinalis NEGAN* promoter (*proNEGAN^C^*) at early corolla developmental stages (0.5-, 1.0- and 2.0-mm in length). The former (B, D and F) showed ridge-associated YFP signals, whereas the latter (C, E and G) had a vein-associated pattern. **(H** and **I)** Confocal fluorescence image at 2.0-mm corolla stage was merged with a bright-field image. The veins -12 to 12 are indicated by white arrowheads. **(J** and **K)** YFP intensity plot within the boxed region in (D) and (E). The relative positions of the veins -7 to 7 are indicated on the x-axis.

To directly test these hypotheses, we generated promoter reporter lines for the *NEGAN^L^* and *NEGAN^C^* alleles in a common genetic background. To this end, we fused the ∼2-kb DNA fragment upstream of the ATG start codon of *NEGAN* with two tandem yellow fluorescent protein (2xYFP) tags, and transformed the *proNEGAN^L^:2xYFP* and *proNEGAN^C^:2xYFP* constructs separately into *M. parishii*, a selfing species that is closely related to *M. lewisii* and *M. cardinalis* and is particularly amenable to stable transformation.^28,38^ Bright-field microscopy images of the unfolded flower buds showed a relatively simple and symmetrical vein system in the tubular portion of the corolla. Each of the five petals has five longitudinal veins. We took the central vein of the ventral petal as a reference point (“0”), and numbered the 12 veins on each side of the reference vein sequentially (**Figures 2H** and **2I**). Confocal microscopy showed that the expression patterns of *proNEGAN^L^:2xYFP* and *proNEGAN^C^:2xYFP* were very similar at the throat of the corolla (the junction between the corolla tube and petal lobes; **Figures 2B-2G**) but markedly different in the tubular portion of the corolla (highlighted by the blue boxes in **Figures 2D** and **2E**). For the *M. lewisii* promoter (*proNEGAN^L^*), YFP signals were concentrated around the nectar guide area (between vein -3 and 3), symmetrically distributed in relation to vein 0 (**Figures 2B, 2D, 2F,** and **2H**). Quantification of YFP signals revealed a “camel-hump” shape with a relatively high and uniform intensity from vein -3 to -1 and vein 1 to 3, corresponding to the two nectar guide ridges, and a slightly lower intensity between the “humps” (vein -1 to 1), corresponding to the “valley” between the two nectar guide ridges (**Figure 2J**). By contrast, the *proNEGAN^C^* signals showed a few discrete, longitudinal gaps in the tubular region of the ventral and lateral petals (**Figures 2C, 2E, 2G,** and **2I**). Quantification of the YFP intensity indicated that these gaps corresponded to the inter-vein regions and the YFP signal peaks coincided with the veins (**Figure 2K**).

We observed that by the corolla reached 2 mm in length, a number of YFP spots became readily visible in the nectar guide region, which were not yet apparent at earlier developmental stages (compare **Figures 2B**, **2D** with **Figure 2F**), suggesting that the *NEGAN* promoters are responsive to the RD process that had kicked in by the 2-mm stage. To rule out the effect of self-activation on the *NEGAN* promoter reporter, we knocked out the native *NEGAN* gene in *M. parishii* via CRISPR/Cas9-mediated genome editing, which resulted in null mutants without anthocyanin spots (**Figures S2A** and **S2B**). We then introduced the *proNEGAN^L^:2xYFP* and *proNEGAN^C^:2xYFP* reporter genes into the *negan*^CR1^ mutant background through crosses. Except without the fluorescent spots, the promoter reporters displayed the same expression patterns in the *negan* null background as in the wild type (**Figures S2C** and **S2D**). These results suggest that the *proNEGAN^L^* and *proNEGAN^C^* reporter patterns observed at early developmental stages (e.g., 1-mm stage) reflect the true prepatterns of *NEGAN^L^* and *NEGAN^C^* expression, which hereafter will be referred to as the “ridge-associated” and “vein-associated”, respectively. Notably, the two different prepatterns are tightly correlated with where pigment production is first initiated in the corresponding species (**Figures 1C** and **1E**), further implicating the critical role of the prepattern of *NEGAN* expression in pigment pattern formation and evolution.

To determine which prepattern represents the ancestral state and which represents the derived state, we generated *NEGAN* promoter reporter lines (also in the *M. parishii* background) for the outgroup species *M. guttatus*, which diverged from the *M. lewisii* species complex ∼20 million years ago (**Figure 1A**).^28^ The spatiotemporal dynamics of pigment spot emergence in *M. guttatus* is very similar to that in *M. lewisii*: first appearing along the two nectar guide ridges and only later emerging between the ridges and spreading to the lower part of the lateral petals (**Figure S3A**). Consistently, confocal fluorescent imaging of the *proNEGAN^G^:2xYFP* reporter showed a “ridge-associated” pattern resembling that of *M. lewisii* (**Figure S3D**). Taken together, these results strongly suggest that in our focal species, the ridge-associated prepattern represents the ancestral state and the vein-associated prepattern evolved later.

These observations also prompted us to investigate *M. parishii*, which is phylogenetically more closely related to *M. cardinalis*^28,39^ but produces dispersed anthocyanin spots in open flowers that are superficially more similar to *M. lewisii* (**Figure 1A**). Intriguingly, examination of the early corolla developmental stages revealed that the spots seem to first emerge along the central veins of the ventral and lateral petals (**Figure S3B**, 3-4 mm stages). To confirm this vein-associated emergence pattern, we transformed *M. parishii* with the *proRTO:CFP-ER* reporter and found that, before the anthocyanin pigments became visible, *RTO* promoter activity was also first detectable in the central veins of the ventral and lateral petals (**Figure S3C**). These results indicate that the *NEGAN* promoter of *M. parishii* may have the vein-associated prepattern. Indeed, confocal fluorescent imaging of the *proNEGAN^P^:2xYFP* reporter revealed the same pattern as observed in *M. cardinalis*, including strong YFP signals along the veins and the longitudinal gaps of signal between the veins (**Figure S3E**). Furthermore, swapping the *NEGAN* alleles between *M. parishii* and *M. cardinalis* in NILs did not cause any conspicuous phenotypic change (**Figures S1B** and **S1C**), corroborating that *NEGAN* alleles in *M. parishii* and *M. cardinalis* have interchangeable function. Based on these results, we conclude that the vein-associated prepattern of *NEGAN* expression has evolved in the common ancestor of *M. parishii* and *M. cardinalis* after splitting from *M. lewisii*.

### Modulation of the promoter strength of the inhibitor *RTO*

Having shown that *M. parishii* and *M. cardinalis* share the same vein-associated prepattern of *NEGAN* expression, next we asked: what accounts for the pattern difference between the two species (i.e., dispersed spots vs. longitudinal stripes)? To test whether the inhibitor *RTO* plays a role, we introgressed the *RTO^C^* allele from *M. cardinalis* into *M. parishii* (**Figure S1D**). The resulting *RTO^C/C^*NIL accumulated conspicuously more anthocyanins in the nectar guide area than the wild-type *M. parishii* (**Figures 3A** and **3B**), suggesting that *RTO* indeed contributes to the pattern difference between *M. parishii* and *M. cardinalis*. Because the anthocyanin phenotype of the *M. parishii RTO^C/C^* NIL was reminiscent of the phenotype observed in some of the previously published *RTO* RNA interference (RNAi) lines in *M. lewisii*,^22^ we speculated that the *M. cardinalis RTO^C^* allele may have weaker expression than the *M. parishii RTO^P^* allele. To test this possibility and to ensure the two alleles experience the same trans-acting environment, we performed RNA sequencing (RNA-seq) analysis of the F_1_ hybrids between the two species. We observed higher allele-specific expression of *RTO^P^* than *RTO^C^* (**Figure 3C**). Consistently, the *M. parishii RTO* promoter showed stronger activation by NEGAN than the *M. cardinalis RTO* promoter in dual-luciferase assays (**Figure 3D**). Taken together, these results suggest that the decrease in *RTO* promoter activity was an important step leading to the *M. cardinalis* pattern.

**Figure 3.**
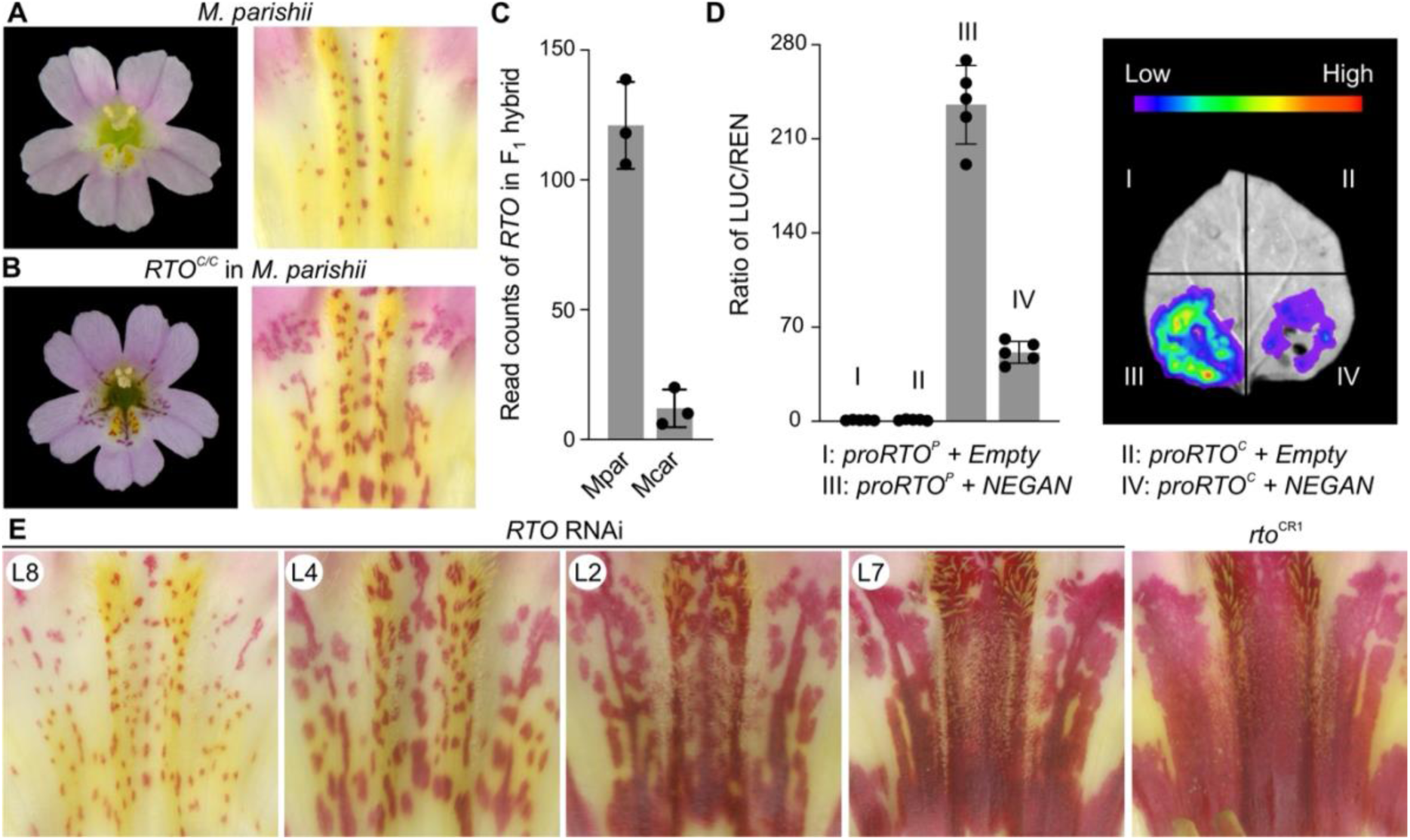
Role of *RTO* promoter activity in pattern variation **(A** and **B)** Front and dissected views of the wild-type *M. parishii* (A) and the *RTO^C/C^* NIL (B). (C) Allele-specific expression analysis of the F_1_ hybrid between *M. parishii* and *M. cardinalis* showed more reads from the F_1_ RNA-seq data were specifically mapped to the *M. parishii* (Mpar) *RTO* allele than the *M. cardinalis* (Mcar) allele. Error bars are 1 SD from three biological replicates. (D) Dual-luciferase assay in tobacco leaves showed the *M. parishii RTO* promoter can be activated much more strongly by NEGAN than the *M. cardinalis* version. Error bars are 1 SD from five biological replicates. A representative live image of the LUC captured in the dark was shown on the right. The *RTO* RNAi and CRISPR (*rto*^CR1^) mutant lines in *M. parishii* display a range of anthocyanin pigment patterns. The edited site for the *rto*^CR1^ is shown in Figure S4.

To explore the phenotypic effect of further reducing *RTO* activity in the *M. parishii* background, we generated a series of *RTO* RNAi lines and CRISPR/Cas9-induced *rto* mutants (**Figures 3E** and **S4**). As expected, the various RNAi lines showed a gradient of phenotypes, with different degrees of anthocyanin expansion. The strongest RNAi lines closely resemble the null mutants. However, none of the mutant or RNAi lines had the discrete, longitudinal stripes found in *M. cardinalis*, indicating there must be other critical factors contributing to the formation of longitudinal stripes.

### Critical role of the bHLH coactivators

To identify the additional components essential for producing the *M. cardinalis* pattern, we took a genetic mapping approach in conjunction with NIL construction in the *M. parishii* background. To this end, we first selected an individual with large, vertically arranged anthocyanin spots from a BC_2_S_1_ population generated by crossing the *M. parishii RTO^C/C^* NIL and *M. cardinalis* and two rounds of backcrossing and selfing (**Figure S5**). Genotyping this individual revealed two large fragments on chromosome 5 (chr5) and chr6 introgressed from *M. cardinalis*. We then backcrossed this individual to the *M. parishii RTO^C/C^* NIL and in the BC_3_S_1_ population, we found that the introgressed chr5 locus produced larger spots and the chr6 locus produced more linearly arranged spots. Individuals homozygous for the *M. cardinalis* allele at both loci formed patterns similar to *M. cardinalis* (**Figure S5**). Further genotyping of this BC_3_S_1_ population delimited both loci to an ∼1 Mb interval. Intriguingly, the chr5 and chr6 interval each contains a subgroup IIIf *bHLH* gene, *ANbHLH3* and *ANbHLH2*,^29^ respectively, that encodes the bHLH component of the anthocyanin-activating MYB-bHLH-WD40 complex in *Mimulus* (hereafter referred to as *BH3* and *BH2* for convenience). This information also allowed us to generate a *BH2^C/C^* and a *BH3^C/C^* NIL in the *M. parishii* background and three combinatorial double NILs with two of the three loci (*RTO*, *BH2*, *BH3*) introgressed as well as a triple NIL with all three loci introgressed (**Figure 4A**).

**Figure 4.**
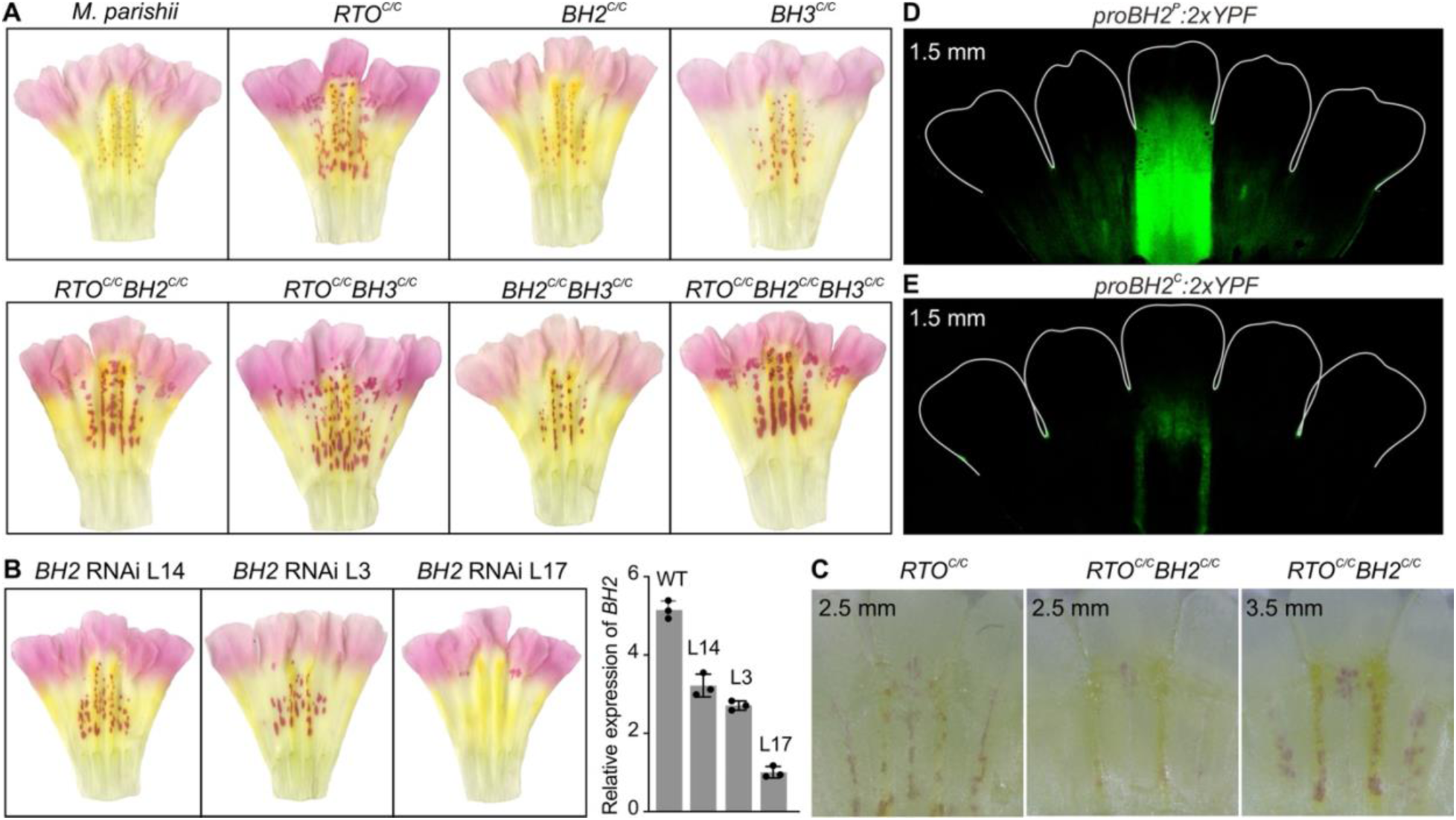
Role of the *BH2* prepattern in the formation of longitudinal stripes (A) Scanned images showing the anthocyanin pigment patterns in various genotypes. (B) *BH2* RNAi lines in the *RTO^C/C^BH3^C/C^* double NIL background. (C) Anthocyanin patterns in the *RTO^C/C^* and *RTO^C/C^BH2^C/C^* NILs at early developmental stages. Spots are readily visible at the 2.5-mm stage in the *RTO^C/C^* NIL, but not until the 3.5-mm stage in the *RTO^C/C^BH2^C/C^* double NIL. **(D** and **E)** The *BH2* promoter reporter line of the *M. parishii* allele (D) showed a broader fluorescence pattern than that of the *M. cardinalis* allele (E).

Given that all NILs carrying the *M. cardinalis BH2^C^* allele showed more linearized pattern than the counterparts carrying the *BH2^P^* allele (**Figure 4A**), we suspected that *BH2* has played a critical role for the spot-to-stripe transition. Consistent with this hypothesis, qRT-PCR showed that *BH2 was* specifically expressed in the tubular portion of the corolla, with a ∼13-fold higher transcript level than that in the petal lobes (**Figure S6A**). Furthermore, knocking down the expression of *BH2* using RNAi in the *RTO^C/C^BH3^C/C^* double NIL background led to reduced or complete lack of spot formation (**Figure 4B**), suggesting that *BH2* is necessary for anthocyanin pattern formation in the corolla tube.

To determine how the *M. cardinalis BH2^C^* allele promotes the linearized arrangement of the spots, we compared the early stages of spot development between the *RTO^C/C^BH2^C/C^* double NIL and the *RTO^C/C^* single NIL. Unexpectedly, we found that anthocyanin spots emerged later in the double NIL than in the single NIL, and became visible on the nectar guide ridges before the central vein of the ventral petal in the *RTO^C/C^BH2^C/C^*double NIL (**Figure 4C**). This observation prompted us to speculate that the *M. cardinalis BH2^C^* allele has a different prepattern of expression than the *M. parishii BH2^P^*allele. To test this idea, we fused the ∼1.5-kb upstream promoter sequence of *BH2* with a 2xYFP tag and transformed the reporter construct into *M. parishii*. Compared to the *M. parishii BH2^P^* promoter (*proBH2^P^:2xYFP*), the *M. cardinalis* version (*proBH2^C^:2xYFP*) showed a much weaker fluorescence intensity (**Figures 4D** and **4E**), which is consistent with its lower allele-specific expression in the F_1_ hybrid (**Figure S6B**). More importantly, while the *proBH2^P^:2xYFP* reporter showed strong fluorescent signals across the nectar guide area (**Figure 4D**), the *M. cardinalis* version had a more discrete distribution, concentrated on the two longitudinal nectar guide ridges (**Figure 4E**). These results implicate a previously unsuspected role of the coactivator prepattern in the diversification of pigment pattern output.

Unlike the *BH2^C/C^* NILs producing more linearized spots, the *BH3^C/C^* NILs showed only larger spots in the wild-type *M. parishii* and the *RTO^C/C^* NIL backgrounds (**Figure 4A**). However, when combined with the *RTO^C^* and *BH2^C^* alleles, the triple NIL (*RTO^C/C^BH2^C/C^BH3^C/C^*) produced a striking pattern with large spots arranged in longitudinal stripes resembling *M. cardinalis* (**Figure 4A**). Consistent with its role in increased spot size, the *BH3^C^* allele had higher expression level than *BH3^P^* in the *M. cardinalis* x *M. parishii* F_1_ hybrid (**Figure 5D**). Furthermore, transforming a construct with the *BH3^P^* coding sequence fused with ∼2.4-kb upstream promoter sequence of *BH3^C^* (*pBH3^C^:BH3 ^P^*) into the *RTO^C/C^BH2^C/C^* NIL recapitulated the striped pattern in transgenic lines (**Figures 5A**). These results confirmed the critical role of *BH3* in forming the longitudinal stripes and suggested that the allelic difference between *BH3^C^* and *BH3^P^* lies in the promoter region.

**Figure 5.**
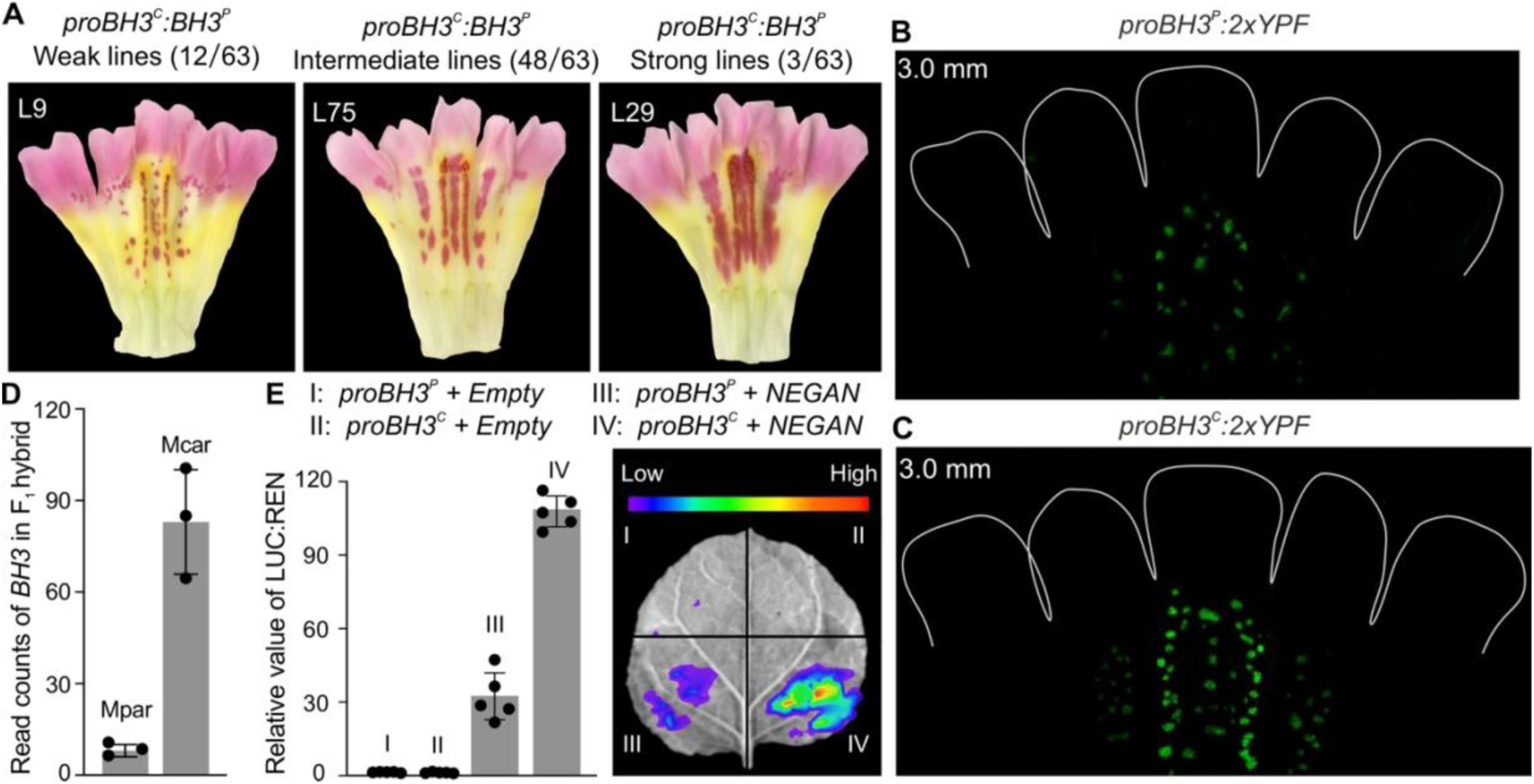
Characterization of the *BH3* promoter activity **(A)** Transgenic phenotype of *proBH3^C^:BH3^P^* in the *RTO^C/C^BH2^C/C^* double NIL background. **(B** and **C)** The *BH3* promoter reporter of the *M. parishii* (B) and *M. cardinalis* alleles (C) showed similar fluorescent patterns coinciding with the anthocyanin pigment spots, with the latter showing stronger signals. (D) Allele-specific expression analysis in the F_1_ hybrid. (E) Dual-luciferase assay in tobacco leaves showing that the *M. cardinalis BH3* promoter is activated more strongly by NEGAN than the *M. parishii* version. A representative live image of the LUC captured in the dark was shown on the right. Error bars are 1 SD from five biological replicates.

To examine the spatial pattern of *BH3* expression, we fused the same ∼2.4-kb upstream promoter sequence of *BH3^P^* and *BH3^C^*, respectively, with a 2xYFP tag, and transformed each reporter construct into *M. parishii*. Surprisingly, we found *BH3* promoter activity coinciding with the pigment spots (**Figures 5B** and **5C**), similar to that of *RTO* (**Figure S3C**). These observations suggested that, just like *NEGAN* and *RTO*, *BH3* is also activated by the MYB-bHLH-WD40 complex. To test this hypothesis, we compared the transcript levels of *BH3*, *NEGAN*, and *RTO* in the wild-type *M. parishii* to that in the *negan* as well as *rto* CRISPR mutants. As expected, qRT-PCR assays showed that all three genes were down-regulated when *NEGAN* was knocked out, and were up-regulated when *RTO* was mutated (**Figure S6C**). In contrast, *BH2* showed no such difference in the mutant lines compared to the wild type. Dual-luciferase assays further confirmed that the MYB-bHLH-WD40 complex can activate the promoter of *BH3^P^* and *BH3^C^*, and the activation is stronger for *BH3^C^* than *BH3^P^* (**Figure 5E**), consistent with the higher transcript level of *BH3^C^* than *BH3^P^*in the F_1_ hybrid (**Figure 5D**).

### Computer simulations using a three-substance RD model

The findings on the importance of the bHLH coactivators in pattern variation confirmed that the previous two-substance, activator-inhibitor model was over-simplified and prompted us to incorporate the coactivator into a new three-substance RD model (**Figure 6A**). In this model, the R2R3-MYB activator (*a*) and the bHLH coactivator (*s*) physically interact, forming a functional activation complex (*c*) that promote pigment production; the R3-MYB inhibitor (*h*) also interact with the bHLH coactivator, removing it from the activation complex. These protein-protein interactions among the three components have been shown in numerous plant systems, including *Mimulus*.^22,34,35^ This three-substance (NEGAN-bHLH-RTO) model has a combination of features of the conventional activator-inhibitor and activator–depleted substrate models, in that the inhibitor is activated by the activation complex but meanwhile inhibits the functionality of the activation complex by depleting the coactivator. Although we have two bHLH paralogs both playing a role in *Mimulus*, their biochemical function as the coactivator is the same from a modeling perspective. Thus, we treated the two bHLH proteins as one substance in the model and we could readily account for their specific differences by adjusting the parameter values. This would also enable more general applicability of the model to other plant systems, since the exact number of bHLH paralogs functioning in pigment pattern formation may differ from species to species.

**Figure 6.**
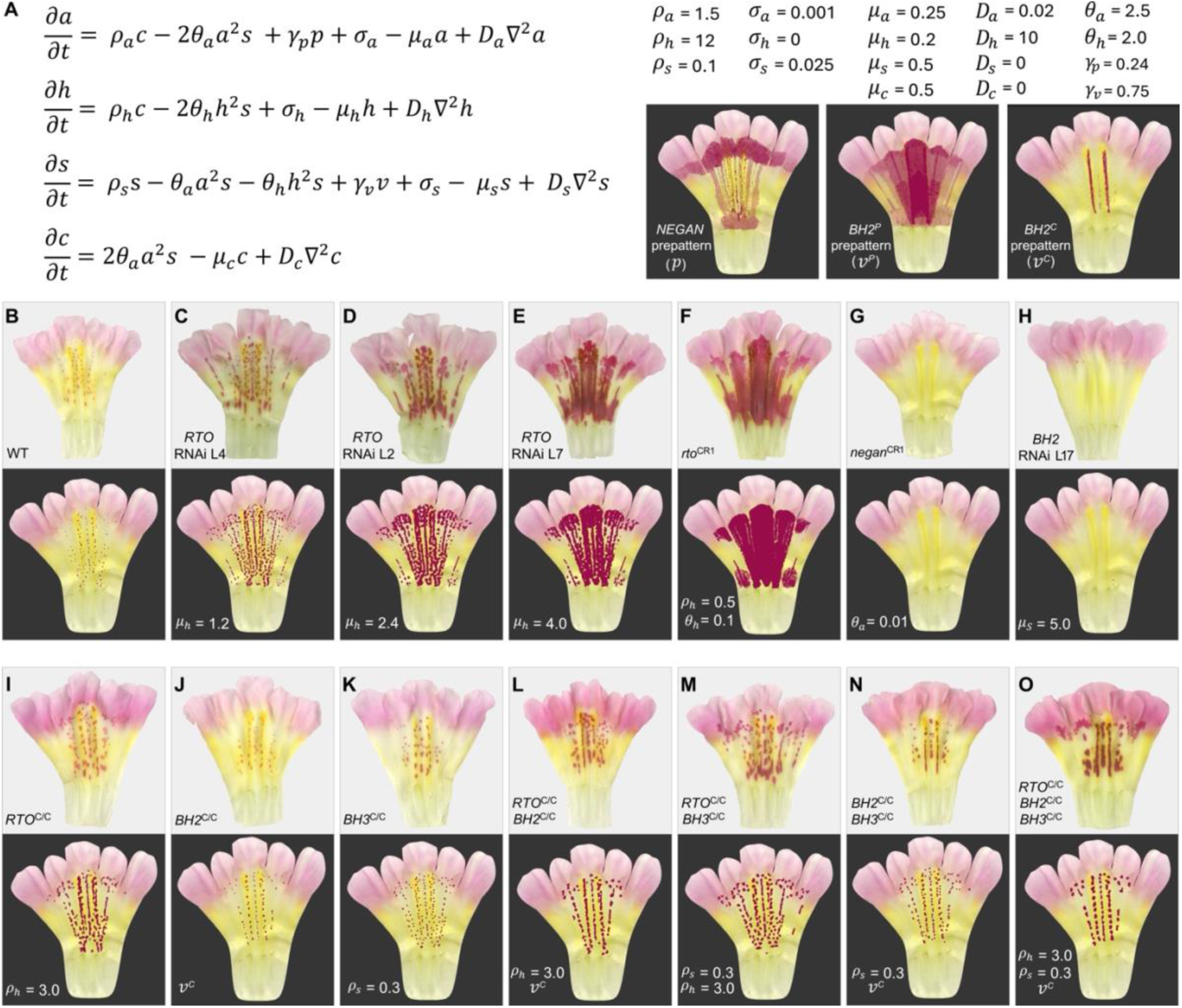
Computer simulation of various pigmentation patterns based on a three-substance RD model. (A) Partial differential equations implemented in the model. 𝑎, ℎ, 𝑠, 𝑐: the concentration of the activator (NEGAN), the inhibitor (RTO), the coactivator (bHLH), and the NEGAN-bHLH activation complex, respectively; 𝜌: activation constant; 𝜃: binding affinity between NEGAN or RTO with bHLH; 𝛾: strength of the prepattern; 𝜎: baseline production rate; 𝜇: degradation rate; 𝐷 : diffusion rate; 𝑝, 𝑣 : the prepattern of NEGAN and bHLH, respectively. Illustrations of the prepatterns and the parameter values used for simulating the wild-type (WT) *M. parishii* pattern are shown on the right of the equations. **(B** to **O)** Simulations of the pigment patterns of the wild-type *M. parishii* (B), various transgenic lines (C-H), and NILs (I-O). The same *NEGAN* prepattern was used for all simulations. The *M. parishii BH2^P^* and *M. cardinalis BH2^C^* prepattern was used as the coactivator prepattern to simulate genotypes that carry the *BH2^P^* and *BH2^C^*allele, respectively. In each panel, the simulated pattern is shown below the real flower image. The difference in parameter values or/and coactivator prepattern from the wild-type conditions is shown in the lower left corner of each panel.

To test whether modifying the RD parameters and the prepatterns in silico can recapitulate the phenotypes generated by experimental perturbations of the corresponding biological entity, we implemented the three-substance model in the program rdpg (reaction- diffusion pattern generator).^23^ We performed computer simulations with a real flower shape as the simulation background and the prepatterns extrapolated from the confocal fluorescent images of the *NEGAN* and *BH2* promoter reporter lines. Our simulations confirmed the large impact of the promoter strength of *RTO* (corresponding to 𝜌_ℎ_ in the model) on pattern variation. Decreasing 𝜌_ℎ_ by 4-fold, a difference in promoter strength between *RTO^P^* and *RTO^C^*indicated by dual-luciferase assay (**Figure 3D**), changed the sparse spots as observed in the wild-type *M. parishii* to irregularly shaped patches that are similar to the *RTO^C/C^* NIL (compare **Figures 6B** and **6I**). Decreasing 𝜌_ℎ_ and 𝜃_ℎ_ simultaneously by ∼20-fold, akin to the CRISPR-mediated knockout of *RTO*, resulted in a pattern closely resembling the *rto*^CR^ phenotype (**Figure 6F)**; and gradually increasing the degradation rate of the inhibitor (𝜇_ℎ_), mimicking the different degrees of *RTO* knockdown by RNAi, also recapitulated the experimental results (**Figures 6C–6E**).

Similarly, either substantially reducing 𝜃_𝑎_ or increasing 𝜇_𝑠_ could completely abolish pigment production as in the *negan* CRISPR mutant or the strong *BH2* RNAi line (**Figures 6G** and **6H**), respectively. Furthermore, replacing the *M. parishii* coactivator prepattern by the *M. cardinalis* version in the simulations led to more linearized spots (**Figures 6J**, **6L**, **6N**, and **6O**) and enhancing the promoter strength of the coactivator (𝜌_𝑠_) generated larger and more rounded spots (**Figures 6K**, **6M**, **6N**, and **6O**). Finally, the combination of decreased 𝜌_ℎ_ (by 4-fold), increased 𝜌_𝑠_(by 3-fold), and the *M. cardinalis* coactivator prepattern, reflecting the genetic difference at the three loci, reproduced the stripy phenotype of the *RTO^C/C^BH2^C/C^BH3^C/C^* triple NIL (**Figures 6O**). Taken together, these simulation results suggest that our three-substance RD model has captured the essence of the RD process underlying the pigment pattern formation in *Mimulus* and confirmed that genetic modulation of the kinetics of the RD system and its prepatterns can cause dramatic pattern variation in nature.

## Discussion

In this study, we have characterized a three-substance RD system underlying the formation of repetitive patterns of pigment spots and stripes in *Mimulus* flowers. Using a combination of genetic analyses, transgenic experiments, and computer simulations, we demonstrate that genetic changes modulating the kinetics of this RD system and its prepatterns are responsible for pigment pattern variation among natural species, from sparsely dispersed spots to longitudinal stripes.

The *Mimulus* RD system includes an R2R3-MYB activator (NEGAN), an R3-MYB inhibitor (RTO), and a coactivator represented by two paralogous bHLH proteins (BH2 and BH3). Although the bHLH coactivator has been long known as an essential component of the MYB-bHLH-WD40 complex that activates anthocyanin biosynthesis in plants,^32–35^ it was not included in the previous two-component RD models to explain *Mimulus* pigmentation patterns, as the coactivator was assumed to remain constant and thus does not contribute to the dynamics of the RD system. It is through genetic mapping of natural pattern variation among species that its role as a critical node of the RD network was revealed. Once the coactivator incorporated, our three-substance model can simultaneously overcome two major limitations of the previous two-substance activator-inhibitor model. First, it no longer assumes an inverse proportion between the rate of activator production and the inhibitor concentration,^3,22,23^ which is inconsistent with the law of mass action^31^ and does not directly correspond to any biochemical entity; instead, the three-substance model is based on experimentally demonstrated protein-protein interactions between the coactivator and the activator or inhibitor, allowing a direct, intuitive biochemical implementation. Second, recent high-throughput mathematical analyses suggested that two-node RD networks may require too restrictive pattern-forming conditions to be biologically realistic. In contrast, RD networks involving three or four substances have much more relaxed pattern-forming conditions and thus are more biologically plausible.^8,40^

The computer simulations also lend support to the validity of this three-substance model. There is no shortage of examples in the literature where computer simulations generated complex pigmentation patterns vividly mimicking real organisms^4,9,23^. However, most of these simulations did not impose biological constrains to the relative parameter values when exploring the parameter space to produce particular patterns, as these relative values are usually unknown. In our simulations, we have imposed several constraints on the relative values based on experimental results. For example, dual luciferase assay suggested that the activation constant of the *M. cardinalis RTO* allele (𝜌_ℎ_) is ∼4 times weaker than the *M. parishii* version, we thus simulated the phenotypic effect of this allele replacement by decreasing 𝜌_ℎ_ 4-fold while keeping all other parameter values the same, with the expectation that the resulting pattern should be similar to the *RTO^C/C^*NIL. The simulation indeed confirmed our expectation (**Figure 6I**). It is reassuring that this model reproduced so many patterns reasonably well (**Figures 6C**- **6O**) by changing one or two parameter values (including prepatterns in some cases) at a time according to experimental results or known biology. A vast number of flowering plants produce similar anthocyanin-based spotty or stripy patterns in the whole or part of their flowers (e.g., butterfly orchid, tiger lily, foxglove, rhododendrons, to name just a few). It is thus possible that this three-substance RD model will be directly applicable to these plant pigmentation systems, and potentially even more widely applicable to patterns based on other pigment types (e.g., carotenoids, betalains, melanins).

This study demonstrates the importance of prepatterns in pattern formation by providing spatial information to the RD process. In *Mimulus*, changes in the prepattern of both the activator (NEGAN) and the coactivator (BH2) are critical to the evolutionary transition from sparsely dispersed spots to longitudinal stripes. The RD model and positional information model represent two of the most prominent theoretical frameworks towards understanding pattern formation,^41,42^ and accumulating evidence suggest that these two mechanisms often act together.^6^ For pigmentation patterns in particular, an emerging theme is that the RD process usually acts downstream of a prepattern: while the former is responsible for generating self-organized, repetitive pattern elements, the latter provides spatial information to specify the location and directionality of the overall pattern.^43–46^ An unresolved issue is how these prepatterns are established in the first place. Addressing this problem in the *Mimulus* system will require elucidation of the regulators and potentially signal transduction pathways upstream of *NEGAN* and *BH2*.

Finally, our work highlights not only the power of RD systems in explaining the rapid diversification of repetitive color patterns in nature, but also the value of natural variation in understanding pattern formation mechanisms. Without genetic mapping of pattern variation among these closely related *Mimulus* species, we would not have recognized the importance of the bHLH coactivator towards building a biologically realistic RD model. Our work on this group of *Mimulus* species has undoubtedly benefited from their short generation time, genetic crossability, and amenability to stable transgenic experiments. However, with the advent of cost-effective chromosome-level genome sequencing and powerful genome-editing tools, genetic and developmental studies that used to be considered infeasible are now becoming achievable in many non-model systems.^47,48^ We expect that more in-depth studies in a wide range of pigmentation systems will likely reveal similar or/and additional types of RD networks that have shaped the tremendous diversity of pigmentation patterns in all multicellular life forms.

## Acknowledgements

We would like to thank Chang Liu, Matt Opel, Cole Geissler, Olivia Delello, and Meghan Moriarty for exemplary plant care at the UConn Botanical Conservatory. This work was supported by an NIH grant R01GM140092 (Y-W.Y. and M.B.); Plant Phenotyping and Imaging Research Centre/Canada First Research Excellence Fund (P.P. and M.C.); Natural Sciences and Engineering Research Council of Canada (NSERC) Discovery Grant 2019-06279 (P.P.); Young Scientist Fostering Funds for the National Key Laboratory for Germplasm Innovation and Utilization of Horticultural Crops (M.L.); and National Excellent Young Scientists (overseas) Fund of National Natural Science Foundation of China (M.L.).

## Author contributions

M.L., P.P., and Y-W.Y. designed the study. Experiments and their analyses were performed by M.L., C.Y., X.Y., N.S., and Y-W.Y. Mathematical model was developed by P.P and M.C. with input from Y-W.Y. Computer simulations were performed by L.R. M.C., M.B., and Y-W.Y. M.L. and Y-W.Y. wrote the manuscript with input from all authors.

## Declaration of competing interests

The authors declare no competing interests.

**Supplemental information** Figures S1–S6 and Table S1, S2 **Materials and Methods**

## Plant materials

The inbred lines used for genetic crosses and tissue collections were as previously described, including *M. guttatus*,^49^ *M. lewisii* LF10,^50^ *M. cardinalis* CE10,^50^ and *M. parishii* Mpar.^38,51^ The self-pollinated *M. parishii* and NILs in the *M. parishii* background were used for transgenic experiments due to its high efficiency in stable transformation.^38^ Plants were grown in the University of Connecticut Botanical Conservatory, with a 16h light and 8h dark photoperiod, and fertilized two or three times a week.

## Genetic crosses and NIL constructions

We used genetic crosses between *M. cardinalis* and *M. lewisii* and between *M. cardinalis* and *M. parishii* to swap their *NEGAN* and *RTO* alleles. The resulting F_1_ was further backcrossed for two rounds and followed by one round of selfing to generate a BC_2_S_1_ population, as illustrated in Figure S1. From this population, the individuals closely resembling the backcrossing parent but with the *NEGAN* or/and *RTO* loci from the other parent were selected as the *NEGAN* or/and *RTO* NILs.

The *RTO^C/C^* NIL in *M. parishii* was refined by four rounds of backcrossing and selfing and was subsequently used for the genetic mapping and the transformation experiment. To determine the size of the introgressed fragment of the *RTO^C/C^* NIL, it was submitted to whole-genome sequencing using the Illumina NovaSeq PE150 platform at Novogene (Sacramento, CA). The Illumina short reads were deposited in National Center for Biotechnology Information (NCBI) (BioProject: PRJNA790954) and were mapped to the *M. parishii* genome (http://mimubase.org/FTP/Genomes/Mparg_v2.0/) using CLC Genomics Workbench 7.5 (QIAGEN, Valencia, CA) to identify homozygous SNPs located in the introgressed fragments from *M. cardinalis*.

To identify the additional loci affecting pigment pattern variation between *M. cardinalis* and *M. parishii*, we generated a BC_2_S_1_ hybrid population by crossing *M. cardinalis* with the *M. parishii RTO^C/C^*NIL through two rounds of backcrossing and selfing. In the BC_2_S_1_ population, one individual with a large, vertically arranged anthocyanin pattern was selected to further backcross with the *M. parishii RTO^C/C^*NIL. The BC_3_ individual was introgressed with two large fragments from *M. cardinalis* on chromosome 5 (between markers MLPC5_350 and MLPC5_7000) and chromosome 6 (between markers MLPC6_150 and MLPC6_5000). The BC_3_ individual was selfed for subsequent genetic mapping and the introgressed fragment of chromosome 5 and chromosome 6 was both shortened to an ∼1 M interval (the genetic mapping process is illustrated in Figure S5).

## RNA extraction, qRT-PCR, and RNA-seq

RNAs used for qRT-PCR analyses or RNA-seq experiments were isolated using the Spectrum Plant Total RNA Kit (STRN250, Sigma-Aldrich). Three biological replicates were used for all qRT-PCR and RNA-seq experiments. For qRT-PCR, 500 ng of total RNAs were used for the cDNA synthesis by GoScript reverse transcriptase (A2791, Promega) and 10 ng generating cDNA were submitted to each qRT-PCR reaction as previously described.^38^ Amplification efficiency for each primer pair was determined using critical threshold values obtained from two fold serial dilutions of pooled cDNA template from 1:4 to 1:64. *MlUBC* was used as a reference gene to normalize expression levels following the delta-delta *C*t method. Primers used for qRT-PCR are listed in Table S1. For RNA-seq, the total RNAs from 5-mm corolla tissues of the *M. parishii* and *M. cardinalis* F_1_ were submitted to RNA library preparation, followed by Illumina Novaseq 6000 platform 150 bp paired-end reads sequencing at Novogene (Sacramento, CA, USA).

## Allele-specific expression analysis

To identify the single nucleotide polymorphism (SNP) variants used for detecting allele-specific expression in the F_1_ hybrid, the genomes of *M. parishii* and *M. cardinalis* were aligned using MUMmer4 v4.0.0.^52^ We extracted SNPs and recalibrated variant quality scores using GATK.^53^ The filtered and biallelic SNPs were used for downstream analysis. Raw reads of the RNA-seq for the *M. parishii* and *M. cardinalis* F_1_ hybrid (NCBI BioProject: PRJNA790954) were processed with Trimmomatic v.0.36^54^ to remove Illumina adaptor sequences and low-quality regions. The resulting clean reads were mapped to the *M. parishii* genome using STAR-WASP^55^ which labels the reads as REF or ALT at the variant sites based on the SNP profile. Reads assigned either REF or ALT genotype were further extracted into separated BAM files. We counted the reads laid in the gene feature for each BAM file using FeatureCounts v2.0.6^56^ and defined it as allele-specific expression.

## Plasmid construction and plant transformation

To study the *RTO* spatial expression pattern on the tubular portion of the corolla among the *M. lewisii* species complex, the *proRTO:CFP-ER* construct was previously transformed into the *M. lewisii* background.^22^ We further transformed the same construct into the *M. parishii* background and then crossed the transgenic with *M. cardinalis* following four rounds of backcrossing to introgress the reporter gene into the *M. cardinalis* background because of the difficulty of the *M. cardinalis* transformation. To examine the spatial expression pattern of the *NEGAN*, *BH2*, and *BH3* genes through two tandem YFP tags, we first amplified a YFP fragment from the pMCS:YFP-GW^57^ and recombined it into pMCS:YFP-GW by LR recombination, resulting in the intermediate vector pMCS:2xYFP. We then amplified the *NEGAN* promoters (2,189-bp upstream of the translation initiation site from *M. guttatus*, 1,958-bp from *M. lewisii*, 2,486-bp from *M. parishii*, and 2,034-bp from *M. cardinalis*), the *BH2* promoters (1,403-bp from *M. parishii* and 1,432-bp from *M. cardinalis*), and the *BH3* promoters (2,648-bp from *M. parishii* and 2,463- bp from *M. cardinalis*). All of the promoter fragments were inserted into the pMCS:2xYFP plasmid and the resulting constructs were transformed into the wild-type *M. parishii*.

To generate the null *negan* and *rto* mutants in *M. parishii*, the endogenous tRNA-processing CRISPR/Cas9 system^58^ was adopted. Because the pRGEB32-BAR vector was originally exploited for rice transformation, we replaced its rice U3 promoter driving sgRNA with the *Arabidopsis thaliana* U6-29 promoter. Two gRNAs were separately designed for the *NEGAN* and *RTO* genes and cloned into the modified pRGEB32-BAR-AtU6.29. The CRISPR/Cas9 constructs were transformed into the wild-type *M. parishii*. To examine the edited sites, the genomic fragments covering the two gRNA sites were amplified in independent transgenic lines. PCR products were ligated into a Zero Blunt TOPO PCR Cloning vector (Thermo Fisher, United States), followed by Sanger sequencing.

The *BH2* RNAi plasmid was generated by cloning a 428-bp *BH2* fragment amplified from the *M. parishii* corolla cDNA into the gateway vector pB7GWIWG2^59^ and was transformed into the *M. parishii RTO^C/C^ BH3^C/C^* double NIL. Given that the *M. lewisii and M. parishii RTO* alleles showed a 99% nucleic acid sequence identity, we transformed the *RTO* RNAi vector constructed previously using *M. lewisii* cDNA into *M. parishii* to generate an *RTO* expression gradient in the *M. parishii* background.

To generate the *proBH3^C^: BH3^P^* rescue plasmid, we first cloned the *BH3* coding sequence from *M. parishii* cDNA into *pMCS:GW*,^57^ resulting the *pMCS:BH3^P^* vector.

Subsequently, we amplified a 2,463-bp fragment upstream of the translation initiation site of the *BH3* gene from *M. cardinalis* genomic DNA and inserted it into *pMCS:BH3^P^*. The resulting *proBH3^C^: BH3^P^* construct was transformed into the *RTO^C/C^BH2^C/C^* NIL in the *M. parishii* background.

All plasmids were transformed into *Agrobacterium tumefaciens* GV3101, followed by plant transformation through the previously described floral spray/infiltration method.^50^ Primers used for plasmid constructions are listed in Table S1.

## Dual-luciferase transient expression

To perform dual-luciferase transient expression in tobacco (*Nicotiana benthamiana*), the full-length coding sequence of *NEGAN* was cloned into the pEarleyGate 202 vector^60^ as effecter construct, which drives the expression of the transgene with the CaMV 35S promoter. Together with the previously generated *35S:BH1* vector,^38^ the *35S:NEGAN* vector was transformed into *Agrobacterium tumefaciens* GV3101. Four DNA fragments upstream of the corresponding translation initiation site, 2,000-bp *RTO^P^* promoter, 2,031-bp *RTO^C^* promoter, 2,648-bp *BH3^P^* promoter, and 2,463-bp *BH3^C^* promoter, were separately inserted into pGreen0800-LUC^61^ to drive the *firefly luciferase* (*LUC*) gene, which also contains the *Renilla* (*REN*) gene for normalization driven by the CaMV 35S promoter. The resulting reporter vectors, *proRTO^P^:LUC:35S:REN*, *proRTO^C^:LUC:35S:REN*, *proBH3^P^:LUC:35S:REN*, and *proBH3^C^:LUC:35S:REN*, were transformed into GV3101 containing the pSoup helper plasmid.

*Agrobacterium* cultures with the corresponding constructs were adjusted to an OD600 of 0.6 with infiltration buffer containing 10 mM MES, 10 mM MgCl_2_ and 150 mM acetosyringone. Each reporter vector was mixed with the *35S:NEGAN* and *35S:BH1* with 1:5:5 (v:v:v) ratio and co-infiltrated into five-week-old leaves using 1-ml needleless syringe. Luciferase images were captured 3 days after infiltration using a NightSHADE LB 985 imaging system (Berthold Technologies, Germany), and the corresponding firefly and renilla luciferase activities were measured with a microplate reader (BioTek Synergy H1, China) using a dual-luciferase reporter assay kit (Promega, USA). Activities of LUC were normalized to REN with the LUC/REN formula and standard errors were calculated from five biological replicates.

## Confocal microscopy

The fluorescence images were captured using an inverted Leica SP8 (Leica, Germany) confocal microscope with HC PL APO CS2 10x/0.4 dry objective. CFP fluorescence for the *RTO* reporter line was taken at 458 nm excitation and 465-600 nm emission, with laser power of 42%, gain value of 703 and pinhole size of 155.3 μm. YFP fluorescence were taken at the 514 nm excitation and 525-600 nm emission. For the *NEGAN* reporter lines, the YFP was imaged with laser power of 35%, gain value of 635 and pinhole size of 106 μm. For the *BH2* reporter lines, the YFP was captured with laser power of 5%, gain value of 717 and pinhole size of 53.1 μm.

For the *BH3* reporter lines, the YFP fluorescence was taken with laser power of 14%, gain value of 888 and pinhole size of 70.7 μm.

## Mathematical modeling and computer simulations

Experimental data from this study and previous studies^22,29^ suggest the following properties of the RD network governing pigmentation patterning in *Mimulus* flowers:

1. Three proteins – the activator NEGAN (*A*), the coactivator bHLH (*S*), and the inhibitor RTO (*H*) – are key components of the RD network;
2. NEGAN, bHLH and another protein, WD40, form a functional activation complex (*C*), which promotes pigment production;^22,29^
3. The activation complex *C* promotes the production of NEGAN and RTO as well as one of the bHLH paralog BH3;
4. RTO can also physically interact with the bHLH coactivator, thereby depleting it from the activation complex *C*.
5. RTO definitely moves between cells, and probably so doe NEGAN;

We account for the spatial inhomogeneity of the generated patterns by considering that the base production of NEGAN and BH2 is modulated by their prepatterns. In this sense, the reaction-diffusion system acts as a translator of larger-scale prepatterns into the detailed patterns of spots, stripes and blotches. And we make two further assumptions based on our current understanding of the system, albeit without experimental evidence yet:

1. WD40 is omnipresent in the flower and consequently it does not play a critical regulatory role in the patterning process.
2. Because the bHLH proteins and the activation complex *C* are large molecules, we assume they do not move between cells in our simulations, although we have built their diffusion into the partial differential equations to allow the versatility and to accommodate future discoveries that may indicate otherwise.

To verify whether these processes can generate pigmentation patterns, we created a computational model (**Figure 6A**) that incorporated the five experimentally verified properties and the two assumptions. For brevity, in the model we denote morphogens NEGAN, bHLH and RTO using uppercase letters *A*, *S* and *H*, respectively (these symbols reflect relations of the proposed model with both the **A**ctivator-**S**ubstrate model and the **A**ctivator-in**H**ibitor model).

Concentrations of morphogens are denoted by the corresponding lowercase letters; the same letters are used as subscripts to identify model parameters pertinent to specific morphogens. The observed pattern is a readout of the concentration of *C*.

For computer simulations, we implemented this set of partial differential equations (**Figure 6A**) in the program rdpg (reaction-diffusion pattern generator).^23^ We input a flower shaped triangular mesh as the simulation domain along with a real flower image as the simulation background, and the prepatterns extrapolated from the confocal fluorescent images of the *NEGAN* and *BH2* promoter reporter lines as an initial condition. Simulated patterns along with their corresponding parameter values are presented in **Figure 6**.

## Notes

### Competing Interest Statement

The authors have declared no competing interest.

## References

1. Turing, A.M. (1952). The chemical basis of morphogenesis. Philos. Trans. R. Soc. Lond. B. Biol. Sci. 237, 37–72. 10.1098/rstb.1952.0012.

2. Gierer, A., and Meinhardt, H. (1972). A theory of biological pattern formation. Kybernetik 12, 30–39.

3. Meinhardt, H. (2012). Turing’s theory of morphogenesis of 1952 and the subsequent discovery of the crucial role of local self-enhancement and long-range inhibition. Interface Focus 2, 407–416. 10.1098/rsfs.2011.0097.

4. Kondo, S., and Miura, T. (2010). Reaction-Diffusion Model as a Framework for Understanding Biological Pattern Formation. Science 329, 1616–1620. 10.1126/science.1179047.

5. Marcon, L., and Sharpe, J. (2012). Turing patterns in development: what about the horse part? Curr. Opin. Genet. Dev. 22, 578–584. 10.1016/j.gde.2012.11.013.

6. Green, J.B.A., and Sharpe, J. (2015). Positional information and reaction-diffusion: two big ideas in developmental biology combine. Development 142, 1203–1211. 10.1242/dev.114991.

7. Hiscock, T.W., and Megason, S.G. (2015). Mathematically guided approaches to distinguish models of periodic patterning. Development 142, 409–419. 10.1242/dev.107441.

8. Scholes, N.S., Schnoerr, D., Isalan, M., and Stumpf, M.P.H. (2019). A Comprehensive Network Atlas Reveals That Turing Patterns Are Common but Not Robust. Cell Syst. 9, 243–257.e4. 10.1016/j.cels.2019.07.007.

9. Kondo, S., and Asai, R. (1995). A reaction–diffusion wave on the skin of the marine angelfish Pomacanthus. Nature 376, 765–768. 10.1038/376765a0.

10. Sick, S., Reinker, S., Timmer, J., and Schlake, T. (2006). WNT and DKK Determine Hair Follicle Spacing Through a Reaction-Diffusion Mechanism. Science 314, 1447–1450. 10.1126/science.1130088.

11. Bouyer, D., Geier, F., Kragler, F., Schnittger, A., Pesch, M., Wester, K., Balkunde, R., Timmer, J., Fleck, C., and Hülskamp, M. (2008). Two-dimensional patterning by a trapping/depletion mechanism: the role of TTG1 and GL3 in Arabidopsis trichome formation. PLoS Biol. 6, e141.

12. Müller, P., Rogers, K.W., Jordan, B.M., Lee, J.S., Robson, D., Ramanathan, S., and Schier, A.F. (2012). Differential Diffusivity of Nodal and Lefty Underlies a Reaction-Diffusion Patterning System. Science 336, 721–724. 10.1126/science.1221920.

13. Menshykau, D., Kraemer, C., and Iber, D. (2012). Branch mode selection during early lung development. PLoS Comput. Biol. 8, e1002377.

14. Economou, A.D., Ohazama, A., Porntaveetus, T., Sharpe, P.T., Kondo, S., Basson, M.A., Gritli-Linde, A., Cobourne, M.T., and Green, J.B. (2012). Periodic stripe formation by a Turing mechanism operating at growth zones in the mammalian palate. Nat. Genet. 44, 348–351.

15. Raspopovic, J., Marcon, L., Russo, L., and Sharpe, J. (2014). Digit patterning is controlled by a Bmp-Sox9-Wnt Turing network modulated by morphogen gradients. Science 345, 566–570. 10.1126/science.1252960.

16. 16. Scacchi, E., Paszkiewicz, G., Thi Nguyen, K., Meda, S., Burian, A., de Back, W., and Timmermans, M.C. (2024). A diffusible small-RNA-based Turing system dynamically coordinates organ polarity. Nat. Plants 10, 412–422.

17. Nakamasu, A., Takahashi, G., Kanbe, A., and Kondo, S. (2009). Interactions between zebrafish pigment cells responsible for the generation of Turing patterns. Proc. Natl. Acad. Sci. 106, 8429–8434.

18. Inaba, M., Yamanaka, H., and Kondo, S. (2012). Pigment pattern formation by contact-dependent depolarization. Science 335, 677. 10.1126/science.1212821.

19. Kaelin, C.B., McGowan, K.A., and Barsh, G.S. (2021). Developmental genetics of color pattern establishment in cats. Nat. Commun. 12, 5127.

20. 20. Connahs, H., Tlili, S., van Creij, J., Loo, T.Y.J., Banerjee, T.D., Saunders, T.E., and Monteiro, A. (2019). Activation of butterfly eyespots by Distal-less is consistent with a reaction-diffusion process. Development 146, dev169367. 10.1242/dev.169367.

21. Davies, K.M., Albert, N.W., and Schwinn, K.E. (2012). From landing lights to mimicry: the molecular regulation of flower colouration and mechanisms for pigmentation patterning. Funct. Plant Biol. 39, 619–638.

22. Ding, B., Patterson, E.L., Holalu, S.V., Li, J., Johnson, G.A., Stanley, L.E., Greenlee, A.B., Peng, F., Bradshaw, H.D., Blinov, M.L., et al. (2020). Two MYB proteins in a self-organizing activator-inhibitor system produce spotted pigmentation patterns. Curr. Biol. 30, 802–814. 10.1016/j.cub.2019.12.067.

23. Ringham, L., Owens, A., Cieslak, M., Harder, L.D., and Prusinkiewicz, P. (2021). Modeling flower pigmentation patterns. ACM Trans. Graph. TOG 40, 1–14.

24. Miyazawa, S., Okamoto, M., and Kondo, S. (2010). Blending of animal colour patterns by hybridization. Nat. Commun. 1, 66. 10.1038/ncomms1071.

25. Patterson, L.B., and Parichy, D.M. (2019). Zebrafish pigment pattern formation: insights into the development and evolution of adult form. Annu. Rev. Genet. 53, 505–530.

26. Irion, U., and Nüsslein-Volhard, C. (2019). The identification of genes involved in the evolution of color patterns in fish. Curr. Opin. Genet. Dev. 57, 31–38.

27. Bieri, E., Rubio, A.O., and Summers, K. (2024). Beyond color and pattern: elucidating the factors associated with intraspecific aggression in the mimic poison frog (Ranitomeya imitator). Evol. Ecol. 38, 621–638. 10.1007/s10682-023-10285-x.

28. Liang, M., Chen, W., LaFountain, A.M., Liu, Y., Peng, F., Xia, R., Bradshaw, H.D., and Yuan, Y.-W. (2023). Taxon-specific, phased siRNAs underlie a speciation locus in monkeyflowers. Science 379, 576–582. 10.1126/science.adf1323.

29. Yuan, Y., Sagawa, J.M., Frost, L., Vela, J.P., and Bradshaw, H.D. (2014). Transcriptional control of floral anthocyanin pigmentation in monkeyflowers (*Mimulus*). New Phytol. 204, 1013–1027. 10.1111/nph.12968.

30. Bar-Yam, Y. (1997). Dynamics of Complex Systems (Addison-Wesley).

31. Yamamoto, L., and Miorandi, D. (2010). Evaluating the robustness of activator-inhibitor models for cluster head computation. In International Conference on Swarm Intelligence (Springer), pp. 143–154.

32. Ramsay, N.A., and Glover, B.J. (2005). MYB–bHLH–WD40 protein complex and the evolution of cellular diversity. Trends Plant Sci. 10, 63–70.

33. Grotewold, E. (2006). The genetics and biochemistry of floral pigments. Annu Rev Plant Biol 57, 761–780.

34. Gonzalez, A., Zhao, M., Leavitt, J.M., and Lloyd, A.M. (2008). Regulation of the anthocyanin biosynthetic pathway by the TTG1/bHLH/Myb transcriptional complex in Arabidopsis seedlings. Plant J. 53, 814–827.

35. Albert, N.W., Davies, K.M., Lewis, D.H., Zhang, H., Montefiori, M., Brendolise, C., Boase, M.R., Ngo, H., Jameson, P.E., and Schwinn, K.E. (2014). A conserved network of transcriptional activators and repressors regulates anthocyanin pigmentation in eudicots. Plant Cell 26, 962–980.

36. LaFountain, A.M., and Yuan, Y.-W. (2021). Repressors of anthocyanin biosynthesis. New Phytol. 231, 933–949. 10.1111/nph.17397.

37. Fishman, L., Stathos, A., Beardsley, P.M., Williams, C.F., and Hill, J.P. (2013). Chromosomal rearrangements and the genetics of reproductive barriers in *Mimulus* (monkeyflowers). Evolution 67, 2547–2560.

38. Liang, M., Foster, C.E., and Yuan, Y.W. (2022). Lost in translation: Molecular basis of reduced flower coloration in a self-pollinated monkeyflower (*Mimulus*) species. Sci. Adv. 8, eabo1113.

39. Nelson, T.C., Stathos, A.M., Vanderpool, D.D., Finseth, F.R., Yuan, Y.W., and Fishman, L. (2021). Ancient and recent introgression shape the evolutionary history of pollinator adaptation and speciation in a model monkeyflower radiation (*Mimulus* section Erythranthe). PLOS Genet. 17, e1009095. 10.1371/journal.pgen.1009095.

40. Marcon, L., Diego, X., Sharpe, J., and Müller, P. (2016). High-throughput mathematical analysis identifies Turing networks for patterning with equally diffusing signals. eLife 5, e14022. 10.7554/eLife.14022.

41. Wolpert, L. (1969). Positional information and the spatial pattern of cellular differentiation. J. Theor. Biol. 25, 1–47.

42. Akam, M. (1989). Making stripes inelegantly. Nature 341, 282–283.

43. Bailleul, R., Curantz, C., Dinh, C.D.-T., Hidalgo, M., Touboul, J., and Manceau, M. (2019). Symmetry breaking in the embryonic skin triggers directional and sequential plumage patterning. PLOS Biol. 17, e3000448. 10.1371/journal.pbio.3000448.

44. Frohnhöfer, H.G., Krauss, J., Maischein, H.-M., and Nüsslein-Volhard, C. (2013). Iridophores and their interactions with other chromatophores are required for stripe formation in zebrafish. Development 140, 2997–3007.

45. Landge, A.N., Jordan, B.M., Diego, X., and Müller, P. (2020). Pattern formation mechanisms of self-organizing reaction-diffusion systems. Dev. Biol. 460, 2–11.

46. Kratochwil, C.F., and Mallarino, R. (2023). Mechanisms underlying the formation and evolution of vertebrate color patterns. Annu. Rev. Genet. 57, 135–156.

47. Andrade, P., Pinho, C., Pérez i de Lanuza, G., Afonso, S., Brejcha, J., Rubin, C.-J., Wallerman, O., Pereira, P., Sabatino, S.J., Bellati, A., et al. (2019). Regulatory changes in pterin and carotenoid genes underlie balanced color polymorphisms in the wall lizard. Proc. Natl. Acad. Sci. 116, 5633–5642.

48. Feiner, N., Brun-Usan, M., Andrade, P., Pranter, R., Park, S., Menke, D.B., Geneva, A.J., and Uller, T. (2022). A single locus regulates a female-limited color pattern polymorphism in a reptile. Sci. Adv. 8, eabm2387.

49. 49. Palm, E., Brady, K., and Van Volkenburgh, E. (2012). Serpentine tolerance in *Mimulus guttatus* does not rely on exclusion of magnesium. Funct. Plant Biol. 39, 679–688.

50. Yuan, Y.W., Sagawa, J.M., Young, R.C., Christensen, B.J., and Bradshaw, H.D. (2013). Genetic dissection of a major anthocyanin QTL contributing to pollinator-mediated reproductive isolation between sister species of *Mimulus*. Genetics 194, 255–263. 10.1534/genetics.112.146852.

51. Fishman, L., Beardsley, P.M., Stathos, A., Williams, C.F., and Hill, J.P. (2015). The genetic architecture of traits associated with the evolution of self-pollination in Mimulus. New Phytol. 205, 907–917.

52. Marçais, G., Delcher, A.L., Phillippy, A.M., Coston, R., Salzberg, S.L., and Zimin, A. (2018). MUMmer4: A fast and versatile genome alignment system. PLOS Comput. Biol. 14, e1005944. 10.1371/journal.pcbi.1005944.

53. McKenna, A., Hanna, M., Banks, E., Sivachenko, A., Cibulskis, K., Kernytsky, A., Garimella, K., Altshuler, D., Gabriel, S., Daly, M., et al. (2010). The Genome Analysis Toolkit: A MapReduce framework for analyzing next-generation DNA sequencing data. Genome Res. 20, 1297–1303. 10.1101/gr.107524.110.

54. Bolger, A.M., Lohse, M., and Usadel, B. (2014). Trimmomatic: a flexible trimmer for Illumina sequence data. Bioinformatics 30, 2114–2120. 10.1093/bioinformatics/btu170.

55. 55. Van De Geijn, B., McVicker, G., Gilad, Y., and Pritchard, J.K. (2015). WASP: allele-specific software for robust molecular quantitative trait locus discovery. Nat. Methods 12, 1061– 1063. 10.1038/nmeth.3582.

56. Liao, Y., Smyth, G.K., and Shi, W. (2014). featureCounts: an efficient general purpose program for assigning sequence reads to genomic features. Bioinformatics 30, 923–930. 10.1093/bioinformatics/btt656.

57. Michniewicz, M., Frick, E.M., and Strader, L.C. (2015). Gateway-compatible tissue-specific vectors for plant transformation. BMC Res. Notes 8, 63. 10.1186/s13104-015-1010-6.

58. Xie, K., Minkenberg, B., and Yang, Y. (2015). Boosting CRISPR/Cas9 multiplex editing capability with the endogenous tRNA-processing system. Proc. Natl. Acad. Sci. 112, 3570–3575. 10.1073/pnas.1420294112.

59. Karimi, M., Inzé, D., and Depicker, A. (2002). GATEWAY^TM^ vectors for Agrobacterium-mediated plant transformation. Trends Plant Sci. 7, 193–195. 10.1016/S1360-1385(02)02251-3.

60. Earley, K.W., Haag, J.R., Pontes, O., Opper, K., Juehne, T., Song, K., and Pikaard, C.S. (2006). Gateway-compatible vectors for plant functional genomics and proteomics. Plant J. 45, 616–629. 10.1111/j.1365-313X.2005.02617.x.

61. Hellens, R.P., Allan, A.C., Friel, E.N., Bolitho, K., Grafton, K., Templeton, M.D., Karunairetnam, S., Gleave, A.P., and Laing, W.A. (2005). Transient expression vectors for functional genomics, quantification of promoter activity and RNA silencing in plants. Plant Methods 1, 1–14. 10.1186/1746-4811-1-13.

