## Supplemental information for "From spots to stripes: Evolution of pigmentation patterns in monkeyflowers via modulation of a reaction-diffusion system and its prepatterns"

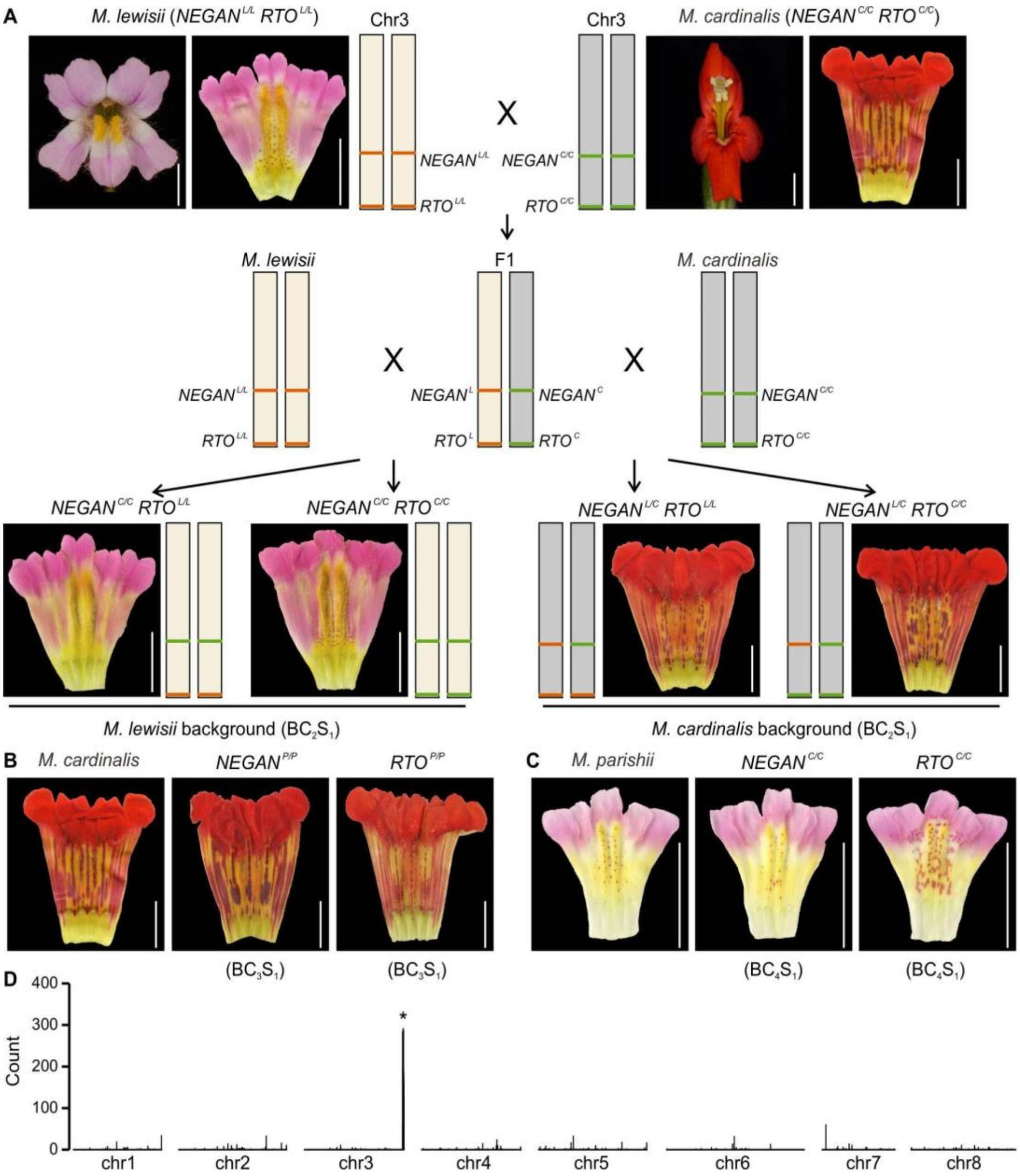


**Figure S1 Allelic swapping of *NEGAN* and *RTO* among *M. lewisii*, *M. cardinalis* and *M. parishii*, related to Figure 2**

**(A)** Cross-design to swap the *NEGAN* and *RTO* alleles between *M. lewisii* and *M. cardinalis* through two rounds of backcrosses and selfing. Only heterozygous *NEGAN*$^{L/C}$ were generated in the *M. cardinalis* background, likely due to segregation distortion known to occur in crosses between *M. lewisii* and *M. cardinalis*.[1]

**(B)** The *NEGAN*$^{P/P}$ and *RTO*$^{P/P}$ NILs in the *M. cardinalis* background were generated through three rounds of backcrosses and selfing.

**(C)** The *NEGAN*$^{C/C}$ and *RTO*$^{C/C}$ NILs in the *M. parishii* background were generated through four rounds of backcrosses and selfing.

**(D)** Whole genome scan of the *RTO*$^{C/C}$ NIL in *M. parishii* for regions that are enriched in homozygous SNPs after aligning the reads to the *M. parishii* reference genome. The peak indicated by the asterisk corresponds to the fragment introgressed from *M. cardinalis*.

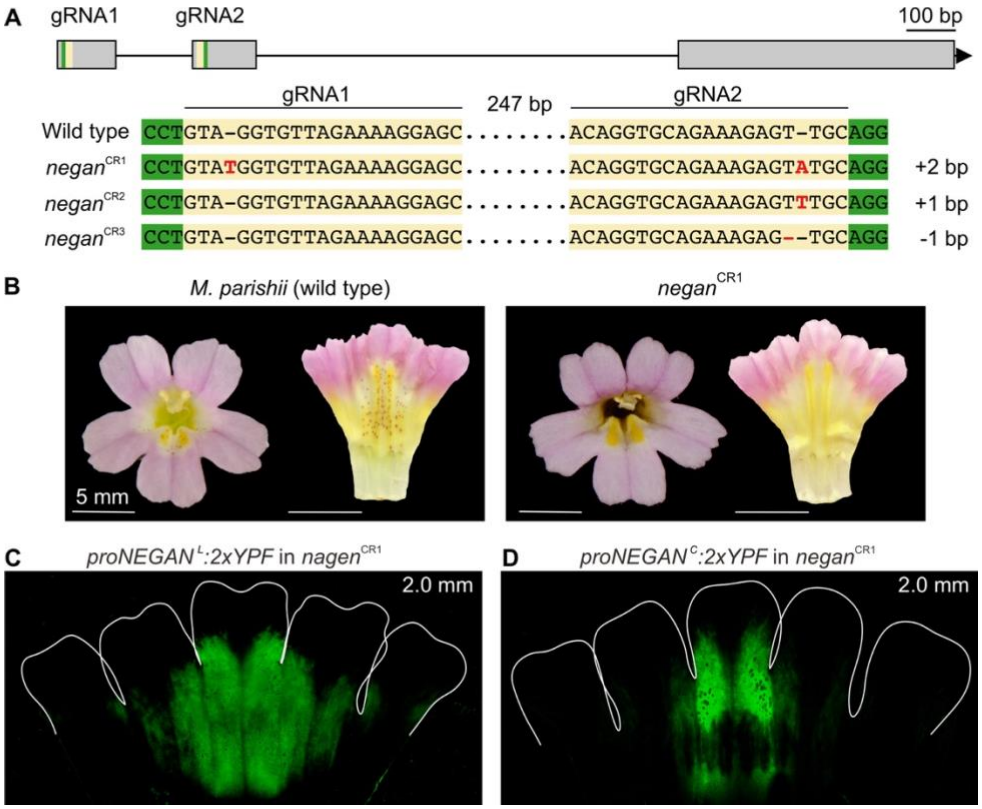


**Figure S2 The YFP fluorescence of the *NEGAN* promoter reflected the true prepatterns of *NEGAN* expression, related to Figure 2**

**(A)** Sanger sequencing of three independent *negan* CRISPR/Cas9 transgenic lines revealed mutations at the two target sites (gRNA1 and gRNA2). The *NEGAN* gene structure is indicated at the top gRNA target sites in yellow and the corresponding PAMs in green. The mutations are shown in red, with the size of deletion (-) or insertion (+) labeled on the right. The sequences between the two gRNAs are represented by the dotted lines.

**(B)** The *negan* CRISPR/Cas9 transgenic line completely lacks anthocyanin spots.

**(C)** The *proNEGAN*$^{L}$ and *proNEGAN*$^{C}$ reporter in the *negan*$^{CR1}$ null mutant showed similar YFP signal patterns to Figures 2F and 2G, respectively, only without the fluorescent spots.

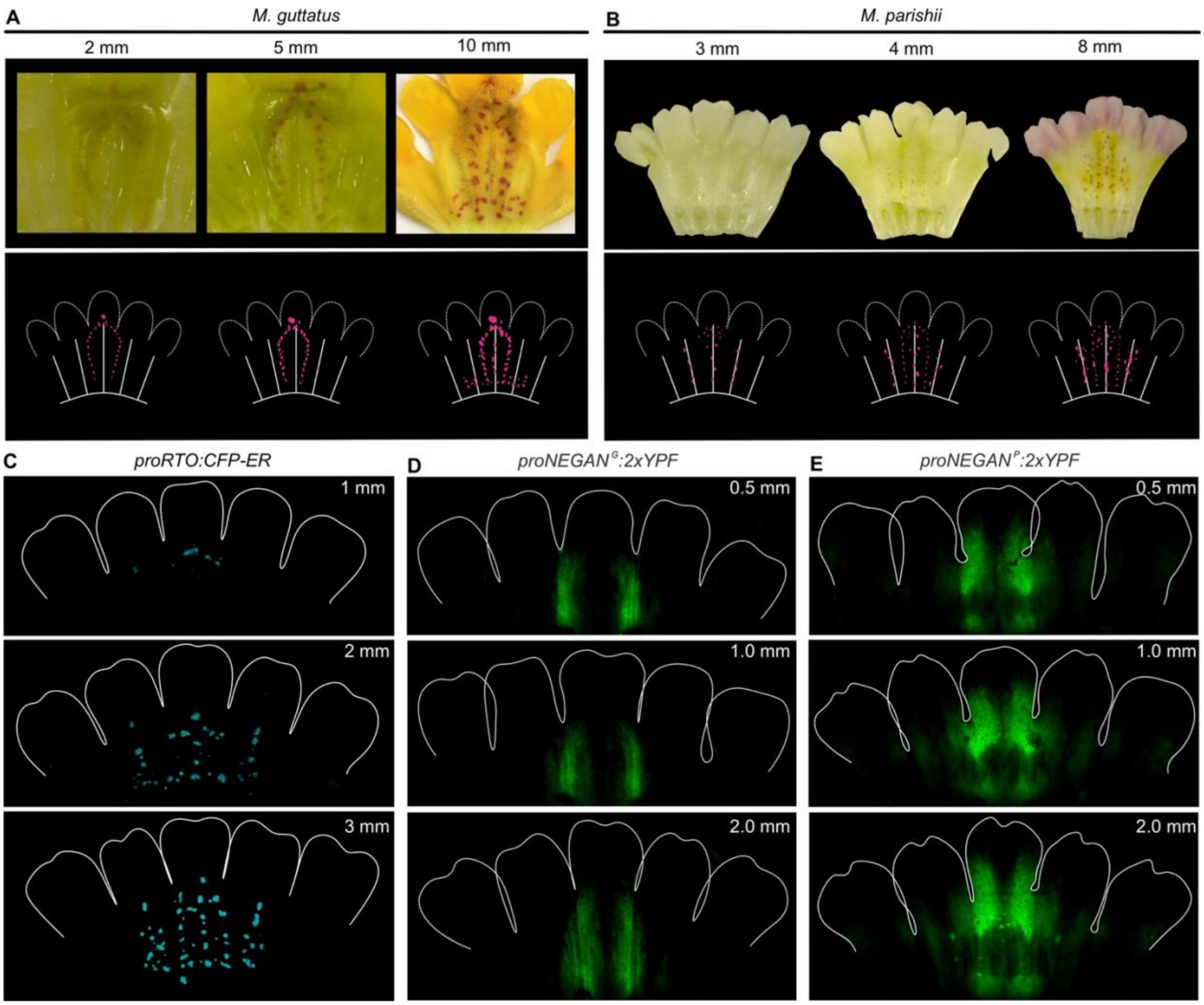


**Figure S3 Pigmentation patterns in *M. guttatus* and *M. parishii*, related to Figures 1 and 2**

**(A** and **B)** The scanned view of *M. guttatus* (A) and *M. parishii* (B) corollas. Corresponding schematic illustrations of the anthocyanin pattern are also shown below the scanned images in (A) and (B).

**(C)** Confocal fluorescent images showing *RTO* promoter activity at early corolla developmental stages (≤ 3 mm) in *M. parishii*.

(**D**) The reporter line of *M. guttatus NEGAN* promoter (*proNEGAN*$^{G}$) showed ridge-associated YFP signals, similar to that of *proNEGAN*$^{L}$ (Figures 2B, 2D and 2F).

(**E**) The reporter line of *M. parishii NEGAN* promoter (*proNEGAN*$^{P}$) showed vein-associated YFP signals, similar to that of *proNEGAN*$^{C}$ (Figures 2C, 2E and 2G).

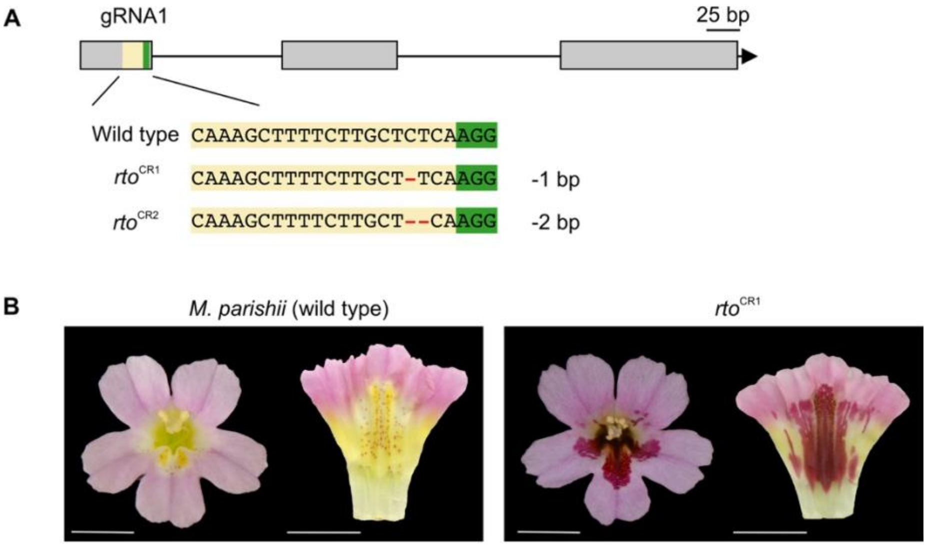


**Figure S4 The flower phenotype of the *RTO* knockout lines, related to Figure 3**
**(A)** Sanger sequencing of two independent *rto* CRISPR/Cas9 transgenic lines revealed deletions at the target sites. The *RTO* gene structure is indicated at the top with gRNA target site in yellow and the corresponding PAM in green.
**(B)** The corolla tube of the *rto* CRISPR/Cas9 knockout line (*rto*$^{CR1}$) accumulated abundant anthocyanin pigments compared to the wild type *M. parishii*. Scale bars are 5 mm.

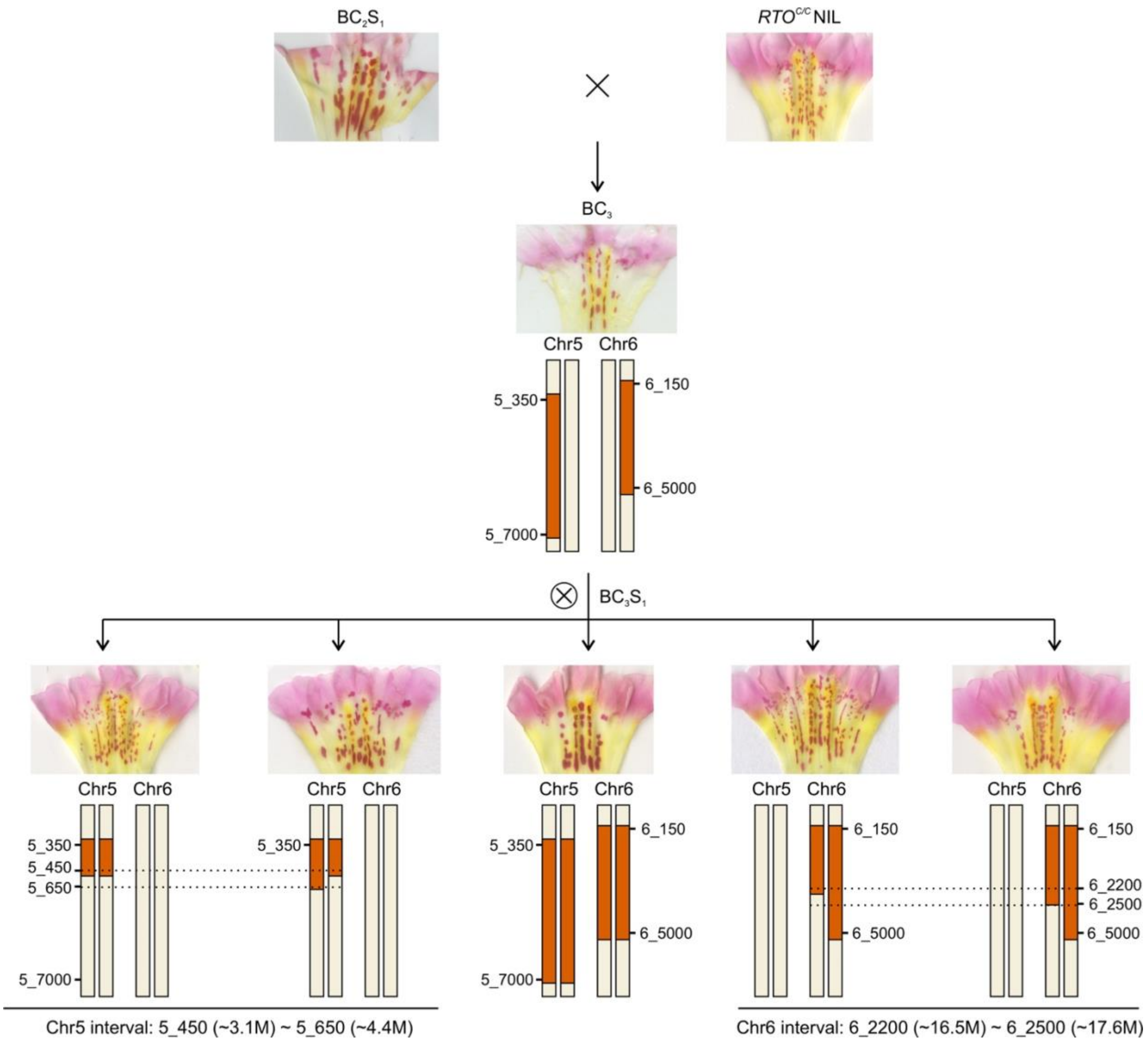


**Figure S5 Fine-scale recombination mapping of the intervals on chromosomes 5 and 6, related to Figure 4**
The *RTO*$^{C/C}$ NIL in the *M. parishiii* background was backcrossed with *M. cardinalis* to identify the additional components involved in the RD model. One $BC_2S_1$ individual with a large, vertically arranged anthocyanin spots was selected for further backcrossing with the *RTO*$^{C/C}$ NIL. For the resulting $BC_3$ individual, two large introgressed fragments between makers MLCP5_350 and MLCP5_7000, and between MLCP6_150 and MLCP6_5000 were identified on each chromosome. The individual with heterozygous chr5 and chr6 intervals was selfed to generate the $BC_3S_1$ population. Genotyping 184 $BC_3S_1$ individuals refined the chr5 and chr6 intervals to ~1 Mb region between MCLP5_450 and MCLP5_650, and between MCLP6_2200 and MCLP6_2500, based on the most informative recombinants. The individual with homologous chr5 and chr6 intervals showed a stripy anthocyanin pattern similar to that of *M. cardinalis*.

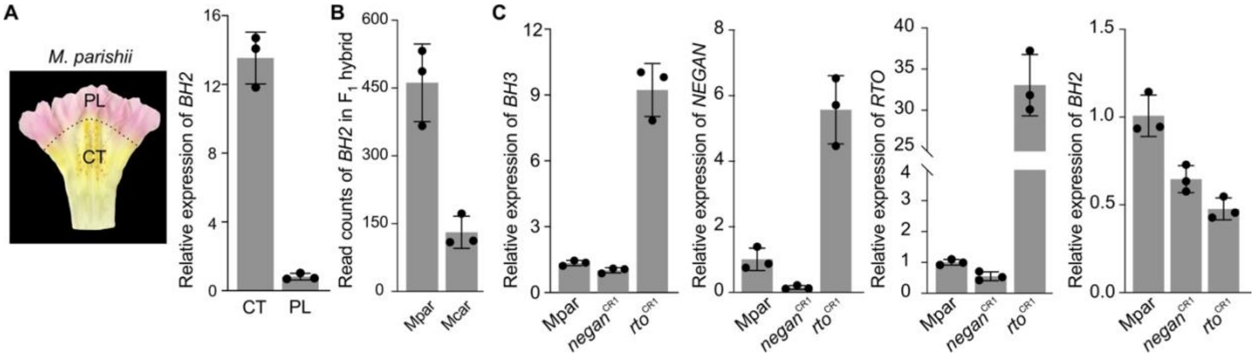


**Figure S6 Relative expression level of *BH2* and *BH3*, related to Figures 4 and 5**

**(A)** *BH2* is preferentially expressed in corolla tube (CT) compared to the petal lobe (PL).

**(B)** Allele-specific expression analysis of the *BH2* gene in the $F_1$ hybrid.

**(C)** Relative transcript level of *NEGAN*, *RTO*, *BH3* and *BH2* in the *M. parashii* wild type (Mpar), the *negan* CRASPR/Cas9 line (*negan*$^{CR1}$) and the *rto* CRASPR/Cas9 line (*rto*$^{CR1}$), as measured by qRT-PCR. The expression of the *BH3*, *NEGAN* and *RTO* genes was downregulated in the *negan*$^{CR1}$ line and upregulated in the *rto*$^{CR1}$ line, indicating NEGAN can activate the transcription of *BH3*, *RTO* and itself. By contrast, BH2 did not show such changes, suggesting *BH2* is not activated by NEGAN.

**Table S1. Primers used for plasmid construction and (q)RT-PCR.**

| Primer name | Sequence (5'-3') | Application |
|---|---|---|
| BP-MpNEGAN-F | GGGGACAAGTTTGTACAAAAAAGCAGGCTGCATGGAAAACACACCTGTAGGTGTTAG | 35S:NEGAN |
| BP-MpNEGAN-R | GGGGACCACTTTGTACAAGAAAGCTGGGTCTTAATTAGGCCCCAGTAGGCC | |
| BP_MpBHLH1_F | GGGGACAAGTTTGTACAAAAAAGCAGGCTGCATGGCTGCTGGAAACCAAGAC | 35S:BH1 |
| BP_MpBHLH1_R | GGGGACCACTTTGTACAAGAAAGCTGGGTCTCCAAATAAGGAACAAGAAAACGTGGGTTC | |
| BP-EYFP-F | GGGGACAAGTTTGTACAAAAAAGCAGGCTTCATGGTGAGCAAGGGCGAGG | pMCS:2xYFP |
| BP-EYFP-R | GGGGACCACTTTGTACAAGAAAGCTGGGTCTCACTTGTACAGCTCGTCCATGC | |
| PMCS-proMgNEGAN_F | AATCTCAGAATTCGCGACGTCATTCATTATATTATGGGTTGATTTACATGG | proNEGAN$^{G}$:2xYFP |
| PMCS-proMgNEGAN_R | CTCGAGGCCTCACGTGTTCGCAGGTGTGTTTTCCTTGCTCTG | |
| PMCS-proMlNEGAN_F | AATCTCAGAATTCGCGACGTTTGCCATAAATCAGTTGTAGAT | proNEGAN$^{L}$:2xYFP |
| PMCS-proMlcpvNEGAN_R | CTCGAGGCCTCACGTGTTCGGGTGTCTTAATTTTCTGCTTCCT | |
| PMCS-proMpNEGAN_F | AATCTCAGAATTCGCGACGTTGCAAACGTGTATAACTGAACTCTG | proNEGAN$^{P}$:2xYFP |
| PMCS-proMlcpvNEGAN_R | CTCGAGGCCTCACGTGTTCGGGTGTCTTAATTTTCTGCTTCCT | |
| PMCS-proMcNEGAN_F | AATCTCAGAATTCGCGACGTTTCCGAACCAACTATTTGCG | proNEGAN$^{C}$:2xYFP |
| PMCS-proMlcpvNEGAN_R | CTCGAGGCCTCACGTGTTCGGGTGTCTTAATTTTCTGCTTCCT | |
| PMCS-proMpBH2-F | AGAATTCGCGACGTCGGTACAACCATTTACTACAAGAAATCAG | proBH2$^{P}$:2xYFP |
| PMCS-proMlcpvBH2-R | TACTCGAGGCCTCACTCCACTATCCACTCTTCACCTCTCTACC | |
| PMCS-proMcBH2-F | AGAATTCGCGACGTCGTATACCTACCAATGTCACCCCTAG | proBH2$^{C}$:2xYFP |
| PMCS-proMlcpvBH2-R | TACTCGAGGCCTCACTCCACTATCCACTCTTCACCTCTCTACC | |
| PMCS-pMpBH3-F-2 | TTTGAAAAATCTCAGCAAGACGACGCCATTGCTAC | proBH3$^{P}$:2xYFP |
| PMCS-pMpBH3-R-2 | GCTCTTATACTCGAGGCGGCTCAACtATTCTTTTAG | |
| PMCS-ProMpcBHLH3-F | TTTGAAAAATCTCAGTCGATTCTGTTACACGTGCCGATC | proBH3$^{C}$:2xYFP |
| PMCS-ProMcBHLH3-R | GCTCTTATACTCGATGTTTTTAGTTTTACCTTCTTTTTTTTGTC | |
| pGreen_proMlcv_RTO_F | TTGATATCGAATTCCTGCAGTTATGCGATGAAATCAAGAACTTGG | proRTO$^{P}$:LUC:35S:REN; proRTO$^{C}$:LUC:35S:REN |
| pGreen_proMlcp_RTO_R | GCTCTAGAACTAGTGGATCCAGTAAAGCTGAGAGTGAAGCTGA | |
| pGreen_proMpBH3-F | TCGACGGTATCGATACAAGACGACGCCATTGCTAC | proBH3$^{P}$:LUC:35S:REN |
| pGreen_proMpBH3-R | CCGCTCTAGAACTAGGGCGGCTCAACtATTCTTTTAG | |
| pGreen_proMcBH3-F | TCGACGGTATCGATATCGATTCTGTTACACGTGCCGATC | proBH3$^{C}$:LUC:35S:REN |

| | | |
|---|---|---|
| pGreen_proMcBH3-R | CCGCTCTAGAACTAGTGTTTTTAGTTTTACCTTCTTTTTTTTGTC | |
| PMCS_proMcBHLH3_F | CCTAGGAGATCTTCGAACACTCGATTCTGTTACACGTGCCGA | proBH3$^{C}$:BH3$^{P}$ |
| PMCS_proMcBHLH3_R | GTACAAACTTGTGATCTCGATGTTTTTAGTTTTACCTTCTTTTTTTTGTC | |
| BP_MpBHLH3_F | GGGGACAAGTTTGTACAAAAAAGCAGGCTGCATGGTTGAGCCGCCGAAGAG | |
| BP_MpBHLH3_R | GGGGACCACTTTGTACAAGAAAGCTGGGTCCCCTTAAATGTAATGAATTGGTTATATTGG | |
| BP_MpRTO_F | GGGGACAAGTTTGTACAAAAAAGCAGGCTGCCAGCTTCACTCTCAGCTTTACT | RTO RNAi |
| BP_MpRTO_R | GGGGACCACTTTGTACAAGAAAGCTGGGTCCAAGCTTGTGCATTCTGCAGA | |
| BP-RNAiMpBH2-F | GGGGACAAGTTTGTACAAAAAAGCAGGCTGCTGTCTGTCAAGTGCTCGATCATGC | BH2 RNAi |
| BP-RNAiMpBH2-R | GGGGACCACTTTGTACAAGAAAGCTGGGTCTTCCACTCTGATTCACCTGCGATC | |
| P1_tRNA-F | AGAGTCGACTAAGAGATTGAACAAAGCACCAGTGGTCTAG | rto CRISPR |
| P1_RTO_MpCDS_g1R | CAAAGCTTTTCTTGCTCTCATGCACCAGCCGGGAATC | |
| P2_RTO_MpCDS_g1F | TGAGAGCAAGAAAAGCTTTGGTTTTAGAGCTAGAAATAGC | |
| P2_RTO_MpCDS_g2R | TATTTCTAGCTCTAAAACTGAGAGCAAGAAAAGCTTTGTTGCACCAGCCGGGAATC | |
| P1_tRNA-F | AGAGTCGACTAAGAGATTGAACAAAGCACCAGTGGTCTAG | negan CRISPR |
| P1_NEGAN_MpCDS_g1R | GTAGGTGTTAGAAAAGGAGCTGCACCAGCCGGGAATC | |
| P2_NEGAN_MpCDS_g1F | GCTCCTTTTCTAACACCTACGTTTTAGAGCTAGAAATAGC | |
| P2_NEGAN_MpCDS_g2R | TATTTCTAGCTCTAAAACGCAACTCTTTCTGCACCTGTTTGCACCAGCCGGGAATC | |
| MLCPV_NEGAN_RTF | AGGAATTTGCTCGAAACTACG | qRT-PCR |
| MLCPV_NEGAN_RTR | CGAGTAATAAATCAGCAAGCCC | |
| MLCPV_RTO_RTF | TGAGTGAGCAAGAACAAGACATC | |
| MLCPV_RTO_RTR | TCAAGAATTATCATGTTTCCTGG | |
| MLCPV_bHLH2_RTF | CTCGGAGCAACTGAACTAGTTCC | |
| MLCPV_bHLH2_RTR | CGAGATCTTCTTCAAGGCTAGCATG | |
| MLCPV_bHLH3_RTF | CTCAAAGACCAGCGCTATCGG | |
| MLCPV_bHLH3_RTR | GGAGACAGGCTACCCACTACTAC | |
| MlUBC | GGCTTGGACTCTGCAGTCTGT | |
| MlUBC | TCTTCGGCATGGCAGCAAGTC | |

Note: the sequences highlighted in red are the adapter sequence necessary for gateway cloning or Gibson assembly.

**Table S2. Key resource table.**

| REAGENT or RESOURCE | SOURCE | IDENTIFIER |
|---|---|---|
| Bacterial and Virus Strains | | |
| Agrobacterium tumefaciens GV3101 | Widely distributed | GV3101 |
| *E. coli* TOP10 competent cells | Widely distributed | TOP10 |
| Biological Samples | | |
| *RTO*$^{C/C}$ NIL in *Mimulus parishii* | This paper | N/A |
| *BH2*$^{C/C}$ NIL in *Mimulus parishii* | This paper | N/A |
| *BH3*$^{C/C}$ NIL in *Mimulus parishii* | This paper | N/A |
| *RTO*$^{C/C}$*BH2*$^{C/C}$ NIL in *Mimulus parishii* | This paper | N/A |
| *RTO*$^{C/C}$*BH3*$^{C/C}$ NIL in *Mimulus parishii* | This paper | N/A |
| *BH2*$^{C/C}$*BH3*$^{C/C}$ NIL in *Mimulus parishii* | | |
| *RTO*$^{C/C}$*BH2*$^{C/C}$*BH3*$^{C/C}$ NIL in *Mimulus parishii* | This paper | N/A |
| *Mimulus cardinalis* × *Mimulus parishii* $BC_2S_1$ seeds | | |
| *Mimulus lewisii* × *Mimulus cardinalis* $BC_2S_1$ seeds | This paper | N/A |
| *RTO*$^{C/C}$ NIL in *Mimulus parishii* × *Mimulus cardinalis* $BC_3S_1$ seeds | This paper | N/A |
| Chemicals, Peptides, and Recombinant Proteins | | |
| D-Luciferin sodium salt | Yeasenbio | Cat#40901 |
| Cetyl Trimethyl Ammonium Bromide (CTAB) | Sigma-Aldrich | Cat#H5882 |
| Critical Commercial Assays | | |
| Dual Luciferase Reporter Assay System | Promega | Cat#E1960 |
| Spectrum Plant Total RNA Kit | Sigma-Aldrich | Cat#STRN250 |

| | | |
|---|---|---|
| GoScript Reverse Transcription system | Promega Corp. | Cat#A5001 |
| SYBR Green master mix | Applied Biosystems | Cat#4367659 |
| Zero Blunt TOPO PCR Cloning Kit | Thermo Fisher Scientific | Cat#450245 |
| GeneArt Gibson Assembly HiFi Master Mix | Thermo Fisher Scientific | Cat#A46628 |
| Gateway BP Clonase II Enzyme mix | Thermo Fisher Scientific | Cat#11789020 |
| Gateway LR Clonase II Enzyme mix | Thermo Fisher Scientific | Cat#11791020 |
| Deposited Data | | |
| RNA sequencing data for *Mimulus cardinalis* × *Mimulus parishii* F1 | NCBI Sequence Read Archive | PRJNA1204532 |
| Whole genome sequencing data for $RTO^{C/C}$ NIL in *Mimulus parishii* | NCBI Sequence Read Archive | |
| *Mimulus parishii* and *Mimulus cardinalis* v2.0 reference genome | Mimubase.org | Mparg_v2.0 and Mcarg_v2.0 |
| Experimental Models: Organisms/Strains | | |
| *Mimulus lewisii* inbred line | Yao-Wu Yuan; available on request | N/A |
| *Mimulus cardinalis* inbred line | Yao-Wu Yuan; available on request | N/A |
| *Mimulus parishii* inbred line | Yao-Wu Yuan; available on request | N/A |
| *Nicotiana benthamiana* | Widely distributed | N/A |
| Oligonucleotides | | |
| Primers used for building transgenic constructs and RT-qPCR, see Table S1 | This paper | N/A |
| Recombinant DNA | | |
| pEarlyGate 202 | *Arabidopsis* Biological Resource Center; described in Earley et al. | CD3-688 |
| pMCS:YFP-GW | *Arabidopsis* Biological Resource Center; described in Marta et al. | CD3-1934 |
| pMCS:GW | *Arabidopsis* Biological Resource Center; described in Marta et al. | CD3-1933 |

| | | |
|---|---|---|
| pB7GWIWG2(II) | https://gatewayvectors.vib.be/collection/pb7gwiwg2ii; described in Karimi et al. | Vector ID: 1_23 |
| pRGEB32-BAR | Addgene; described in Charles. | 126072 |
| pGreen0800-LUC | described in Roger et al. | N/A |
| proRTOP:CFP-ER | described in Ding et al. | N/A |
| 35S:NEGAN (pEarlyGate 202 plasmid) | This paper | N/A |
| 35S:BH1 (pEarlyGate 202 plasmid) | This paper | N/A |
| proRTO$^{P}$:LUC:35S:REN (pGreen0800-LUC plasmid) | This paper | N/A |
| proRTO$^{C}$:LUC:35S:REN (pGreen0800-LUC plasmid) | This paper | N/A |
| proBH3$^{P}$:LUC:35S:REN (pGreen0800-LUC plasmid) | This paper | N/A |
| proBH3$^{C}$:LUC:35S:REN (pGreen0800-LUC plasmid) | This paper | N/A |
| proNEGAN$^{G}$:2xYFP (pMCS:YFP plasmid) | This paper | N/A |
| proNEGAN$^{L}$:2xYFP (pMCS:YFP plasmid) | This paper | N/A |
| proNEGAN$^{C}$:2xYFP (pMCS:YFP plasmid) | This paper | N/A |
| proNEGAN$^{P}$:2xYFP (pMCS:YFP plasmid) | This paper | N/A |
| proBH2$^{C}$:2xYFP (pMCS:YFP plasmid) | This paper | N/A |
| proBH2$^{P}$:2xYFP (pMCS:YFP plasmid) | This paper | N/A |
| proBH3$^{C}$:2xYFP (pMCS:YFP plasmid) | This paper | N/A |
| proBH3$^{P}$:2xYFP (pMCS:YFP plasmid) | This paper | N/A |
| rto crispr (pRGEB32-BAR plasmid) | This paper | N/A |
| negan crispr (pRGEB32-BAR plasmid) | This paper | N/A |

| | | |
|---|---|---|
| BH2 RNAi<br>(pB7GWIWG2 plasmid) | This paper | N/A |
| proBH3$^{C}$: BH3$^{P}$<br>(pMCS:GW plasmid) | This paper | N/A |
| Software and Algorithms | | |
| ImageJ | NIH; https://ImageJ.nih.gov/ij/ | N/A |
| CLC Genomics<br>Workbench 7.5 | QIAGEN | N/A |
| MUMmer4 v4.0.0. | Described in Guillaume et al. | N/A |
| GATK | Described in Aaron et al. | N/A |
| Trimmomatic v.0.36 | Described in Anthony et al. | N/A |
| STAR-WASP | Described in Bryce et al. | N/A |
| FeatureCounts v2.0.6 | Described in Yang et al. | N/A |
